# Explosive seed dispersal depends on SPL7 to ensure sufficient copper for localized lignin deposition via laccases

**DOI:** 10.1101/2022.04.21.488998

**Authors:** Miguel Pérez-Antón, Ilsa Schneider, Patrizia Kroll, Hugo Hofhuis, Sabine Metzger, Markus Pauly, Angela Hay

**Author notes:** Angela Hay, MPIPZ, Carl-von-Linné-Weg 10, 50829 Cologne, Germany., **Email:**. **Author Contributions** A.H. and M.P-A. designed research; M.P-A., I.S., P.K., H.H. performed research; M.P. and S.M. contributed lignin and copper measurements, respectively; M.P-A. analyzed data; A.H. and M.P-A. wrote the article.

## Abstract

Exploding seed pods evolved in the Arabidopsis relative, *Cardamine hirsuta*, via morphomechanical innovations that allow the storage and rapid release of elastic energy. Asymmetric lignin deposition within endocarp*b* cell walls is one such innovation that is required for explosive seed dispersal and evolved in association with the trait. However, the genetic control of this novel lignin pattern is unknown. Here, we identify three lignin-polymerizing laccases, LAC4, 11 and 17, that precisely co-localize with, and are redundantly required for, asymmetric lignification of endocarp*b* cells. By screening for *C. hirsuta* mutants with less lignified fruit valves, we found that loss of function of the transcription factor gene *SQUAMOSA PROMOTER BINDING PROTEIN-LIKE 7* (*SPL7*) caused a reduction in endocarp*b* cell wall lignification and a consequent reduction in seed dispersal range. SPL7 is a conserved regulator of copper homeostasis and is both necessary and sufficient for copper to accumulate in the fruit. Laccases are copper-requiring enzymes. We discovered that laccase activity in endocarp*b* cell walls depends on the SPL7 pathway to acclimate to copper deficiency and provide sufficient copper for lignin polymerization. Hence, SPL7 links mineral nutrition to efficient dispersal of the next generation.

## Introduction

Exploding seed pods are one of many different adaptations that plants have evolved to disperse their seeds. This huge diversity reflects the important ecological and evolutionary consequences of dispersal, including the ability to change or expand a species’ range (1, 2). *Cardamine hirsuta* is a small, ruderal weed, related to the plant model *Arabidopsis thaliana* (3). Unlike *A. thaliana*, it uses an explosive mechanism to disperse its seeds. In the seed pods of *C. hirsuta*, two exploding valves coil back rapidly, firing the seeds at speeds greater than 10 m s^-1^ to disperse over a large area (4). This mechanism requires localized lignin deposition in a single cell layer of the fruit valve called the endocarp*b* (end*b*), as mutant fruit that lack end*b* cells fail to explode (4). A striking feature of end*b* cells in *C. hirsuta* is the asymmetric deposition of lignin in three stiff rods connected by thin hinges (4). End*b* cell walls are uniformly lignified in *A. thaliana* and other species with non-explosive fruit in the Brassicaceae family (5). Explosive seed dispersal evolved once in this family, in the *Cardamine* genus, and asymmetric lignin deposition is strictly associated with this trait in *Cardamine* species (4). Currently, the genetic basis of this localized lignin pattern is unknown.

Asymmetric lignin deposition is known to play a key role in the mechanics of exploding seed pods. Simulations of a mathematical model that described the elastic energy in the fruit valve were compared using an asymmetrically hinged versus a uniformly lignified wall geometry. These results showed that the hinged wall geometry is critical for the explosive release of stored elastic potential energy, allowing the fruit valve to employ a rapid release mechanism like a toy slap bracelet (4). Predictions from these model simulations were tested genetically, by creating transgenic plants with uniformly lignified end*b* cells. These seed pods failed to explode, showing that the asymmetric pattern of lignin deposition in end*b* cells is required for explosive seed dispersal (4).

Lignin is a polymer that imparts stiffness and hydrophobicity to the plant cell wall. It is composed of monolignols, which are synthesized from phenylalanine in the cytosol and exported to the cell wall where they are activated by oxidation (6). Laccases (LAC) and type III peroxidases (PER) are two different types of secreted enzymes that catalyse this oxidation. The lignin polymer is then formed by non-enzymatic random coupling of activated monolignols. While the biosynthesis of monolignols is well understood, it is less clear how localized patterns of lignin deposition are produced. Targeted export of monolignols to specific cell wall domains is unlikely to contribute to lignin patterning, because monolignols are highly mobile in the apoplast, and do not perturb lignin patterning when applied exogenously (7-9). In contrast to this, LAC and PER enzymes often co-localize precisely with subcellular patterns of lignin (10-12). Type III PER and LAC enzymes are encoded by large gene families with 73 and 17 members, respectively, in *A. thaliana* (13-15). Genetic and biocatalytic redundancy within and between these large enzyme families has made it difficult to ascribe biological functions to individual genes. Although a mixed set of oxidative enzymes are likely to contribute to lignification in different cells and tissues (10), the relative requirement for LACs versus PERs tends to differ between cell types. For example, lignification of Casparian strips in the root endodermis is PER-rather than LAC-dependent, as a quintuple PER mutant (*per3 9 39 72 64*) abolished Casparian strips, while a nonuple LAC mutant (*lac1 3 5 7 8 9 12 13 16*) had no effect (16). By contrast, the lignification of xylem tissues depends on laccases. Lignin content was reduced in the stems of *lac4 17* double mutants and further reduced in *lac4 11 17* triple mutants, which failed to grow past the seedling stage (17, 18).

LACs and PERs are secreted glycoproteins that require metal co-factors for enzymatic activity. Type III PERs are heme-containing proteins that reduce H_2_O_2_ to oxidize monolignols. For this reason, PER-dependent lignification requires reactive oxygen species (ROS) produced by NADPH oxidases (respiratory burst oxidase homologs) (12, 19). On the other hand, LACs are multi-copper proteins that, unlike PERs, reduce O_2_ to H_2_O in order to oxidize monolignols, and do not require ROS. Laccases bind four Cu ions: one Cu ion participates in the oxidation of the substrate and a cluster of three participate in the reduction of O_2_ to H_2_O (20). The cycling of Cu between an oxidized (Cu^2+^) and a reduced state (Cu^1+^) is used in many different redox reactions and electron transport. Thus, Cu is an essential micronutrient in nearly all eukaryotic organisms. However, when Cu ions are present in excess, redox cycling can catalyze the production of highly toxic ROS and cause cellular damage. Therefore, Cu homeostasis is tightly controlled in plants (21).

Plants take up Cu from the soil, and the bioavailability of Cu varies depending on soil type, though natural soils with high Cu levels are rare (22). To acclimate to Cu deficiency, *A. thaliana* regulates Cu homeostasis via the SQUAMOSA PROMOTER-BINDING PROTEIN-LIKE 7 (SPL7) transcription factor. SPL7 is one of 16 SPL proteins in *A. thaliana*, which all bind DNA sequences with a core GTAC motif (23). The majority of this gene family are targeted for post-transcriptional degradation by miR156/157, but *SPL7* is not (24). SPL7 is homologous to COPPER RESPONSE REGULATOR 1 (CRR1) in the green alga *Chlamydomonas reinhardtii* (25, 26). Under copper limiting conditions, this evolutionarily conserved switch, activates the transcription of genes that increase Cu uptake and economize on the use of available Cu (27-29). In this way, the SPL7 pathway ensures that when Cu is limiting, sufficient Cu is available for the function of essential cuproproteins such as plastocyanin, which is required for photosynthetic electron transfer and cannot be replaced (30).

Here, we describe the genetic control of localized lignin deposition by three multi-Cu laccases in end*b* cell walls of explosive fruit. LAC4, 11 and 17 precisely co-localize with, and are required for, the unique pattern of lignin in these cells. The non-lignified end*b* cell walls found in *lac4 11 17* triple mutants were phenocopied by loss of *SPL7* function. Our findings show that laccase activity in end*b* cell walls requires SPL7, in order to acclimate to Cu deficiency and provide sufficient Cu to polymerize lignin.

## Results

### *SPL7* is required for end*b* lignification

To identify genes required for localized lignin deposition in end*b* cells, we performed a forward genetics screen in *C. hirsuta* for mutants with less lignified fruit valves. The *less lignin 1* (*lig1*) mutant showed reduced lignification of end*b* secondary cell walls (SCWs) (Fig. 1A-C). The phenotype was particularly pronounced in end*b* cells compared to other lignified cell types in the fruit (Fig. 1B-C). By quantifying acetylbromide soluble lignin, we found that lignin content was significantly reduced in the fruit valves of *lig1* compared to wild type (Fig. 1D). In addition, mature *lig1* fruit buckled along their edge, compared to the straight edge of wild-type fruit (Fig. 1A, D). The timing of this buckling coincided with the lignification of end*b* cells, suggesting it may be a consequence of reduced lignin. We also found a significant reduction in the seed dispersal range of *lig1* plants (Fig. 1 E). Maximum dispersal distance was reduced by 0.5 m in *lig1* and significantly fewer *lig1* than wild-type seeds were dispersed further than 0.75 m (Fig. 1 E). Therefore, lignification of valve end*b* SCWs is critical for explosive seed dispersal in *C. hirsuta*,

**Figure 1.**
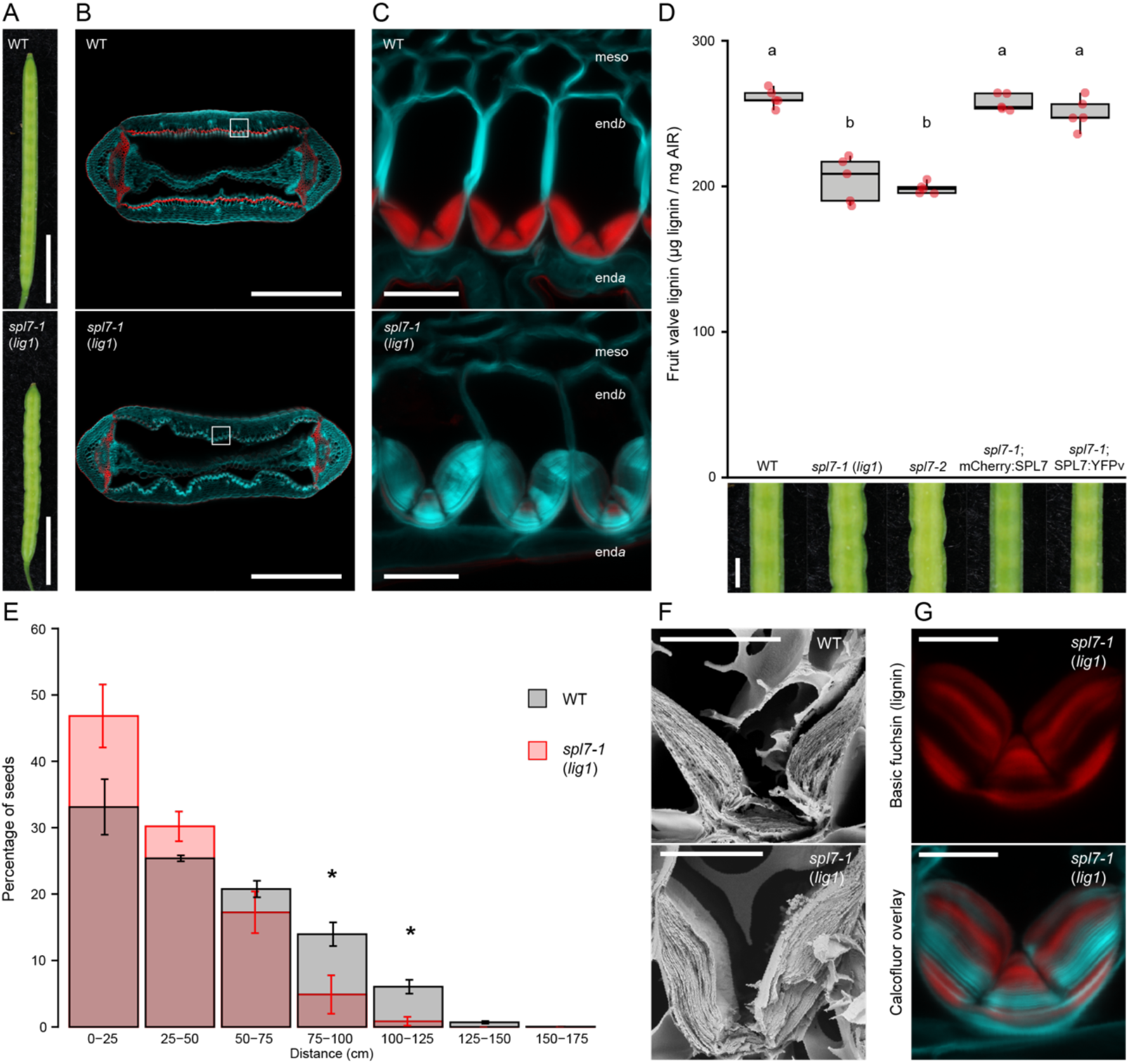
*SPL7* is required for end*b* lignification in *C. hirsuta* fruit. (*A*) Mature wild-type and *spl7-1* (*lig1*) fruit. (*B-C*) Lignin, stained red with basic fuchsin, in cross sections of wild-type and *spl7-1* (*lig1*) fruit (*B*) and end*b* cells (*C*), cell walls stained cyan with calcofluor white. Cells in (*C*) correspond to regions of the valve indicated by white boxes in (*B*). (*D*) Boxplot of lignin concentration, shown as µg of acetyl bromide soluble lignin per mg of alcohol-insoluble residue (AIR), in mature fruit valves, and close-up view of fruit margin, in wild type, *spl7-1* (*lig1*), *spl7-2, spl7-1; mCherry:SPL7* and *spl7-1; SPL7:YFPv*. Plot shows median (thick black line), n = 5 biological replicates per genotype (red dots) where each replicate contains 16 valves from two plants, different letters denote statistical significance at *P* < 0.05 using Kruskal-Wallis and Fisher’s least significant difference as post hoc analysis. (*E*) Barplot of seed dispersal in wild type (grey) and *spl7-1* (red), showing the percentage of seeds in each distance bin. Error indicates SEM, n = 10039 total seeds dispersed by 4 plants per genotype, * denotes statistical significance at *P* < 0.05 using Student’s *t*-test. (*F*) Scanning electron micrographs of wild-type and *spl7-1* cryo-fractured fruit showing end*b* SCW layers. (*G*) Variable lignin deposition (stained red with basic fuchsin) in *spl7-1* end*b* cells, overlaid with calcofluor white cell wall stain (cyan). Confocal micrographs show z-axis sum projections of transverse fruit sections (*B, C, G*). All plants were supplemented with 0.5 mM CuSO_4_ to ensure plant growth and fruit development of *spl7* mutants. Abbreviations: meso: mesocarp, end*b*: endocarp*b*, end*a*: endocarp*a*. Scale bars: 5 mm (*A*), 500 µm (*B*), 20 µm (*C*), 1 mm (*D*), 10 µm (*F-G*).

We used mapping by sequencing to identify the causal mutation of *lig1* as a C>T mutation that introduced a P443S substitution in the *C. hirsuta* ortholog of the transcription factor SQUAMOSA PROMOTER BINDING PROTEIN-LIKE 7 (SPL7, CARHR194170) (Fig. S1). To verify that loss of *SPL7* function caused a similar phenotype to the recessive *lig1* allele, we used CRISPR/Cas9 to introduce a single nucleotide deletion that resulted in a truncated 66 amino acid protein lacking the conserved SBP DNA-binding domain (Fig. S1). These *spl7* mutant fruit valves had reduced lignin content and less lignified end*b* SCWs, indistinguishable from *lig1* (Fig. 1D, Fig. S1). Moreover, an allelism test showed that *lig1* is a *spl7* allele and likely to represent complete loss of *SPL7* function in *C. hirsuta* (Fig. S1). We showed that all *lig1* phenotypes were fully complemented by expressing the wild-type *SPL7* genomic locus, tagged at either the N-terminus with mCherry (*pSPL7::mCherry:SPL7*) or the C-terminus with YFP (*pSPL7::SPL7:YFP*) (Fig. 1D, Fig. S1). Moreover, a 253 aa truncated SPL7 protein, including the SBP DNA-binding domain, NLS and transcriptional activation domain (*pSPL7::SBP:GFP*), was sufficient to fully rescue the fruit defects of *lig1* to wild type (Fig. S1). A similar finding was reported for *A. thaliana* SPL7 (31) and suggests that lignification of end*b* SCWs requires transcriptional responses that are regulated by SPL7. Based on these findings, we renamed *lig1* as *C. hirsuta spl7-1* and the CRISPR/Cas9 allele *spl7-2*.

To investigate the localization of SPL7 in *C. hirsuta* fruit during end*b* SCW formation, we analyzed transcriptional *SPL7* reporters and functional SPL7 protein fusions. *pSPL7::GFPnls* localized to end*b* cells before and during the deposition of lignified SCWs in *C. hirsuta* fruit valves (Fig. 2A-B). We observed an identical pattern in complementing *pSPL7::SBP:GFP* lines (Fig. S2). We could detect *SPL7* transcripts in most *C. hirsuta* plant tissues (Fig. S2). However, we could not reliably detect the tagged full-length SPL7 proteins that complemented *C. hirsuta spl7-1*, similar to previous reports for *A. thaliana* SPL7 (31). In summary, SPL7 accumulates in end*b* cells where the lignification defect is observed in *spl7* mutants.

**Figure 2.**
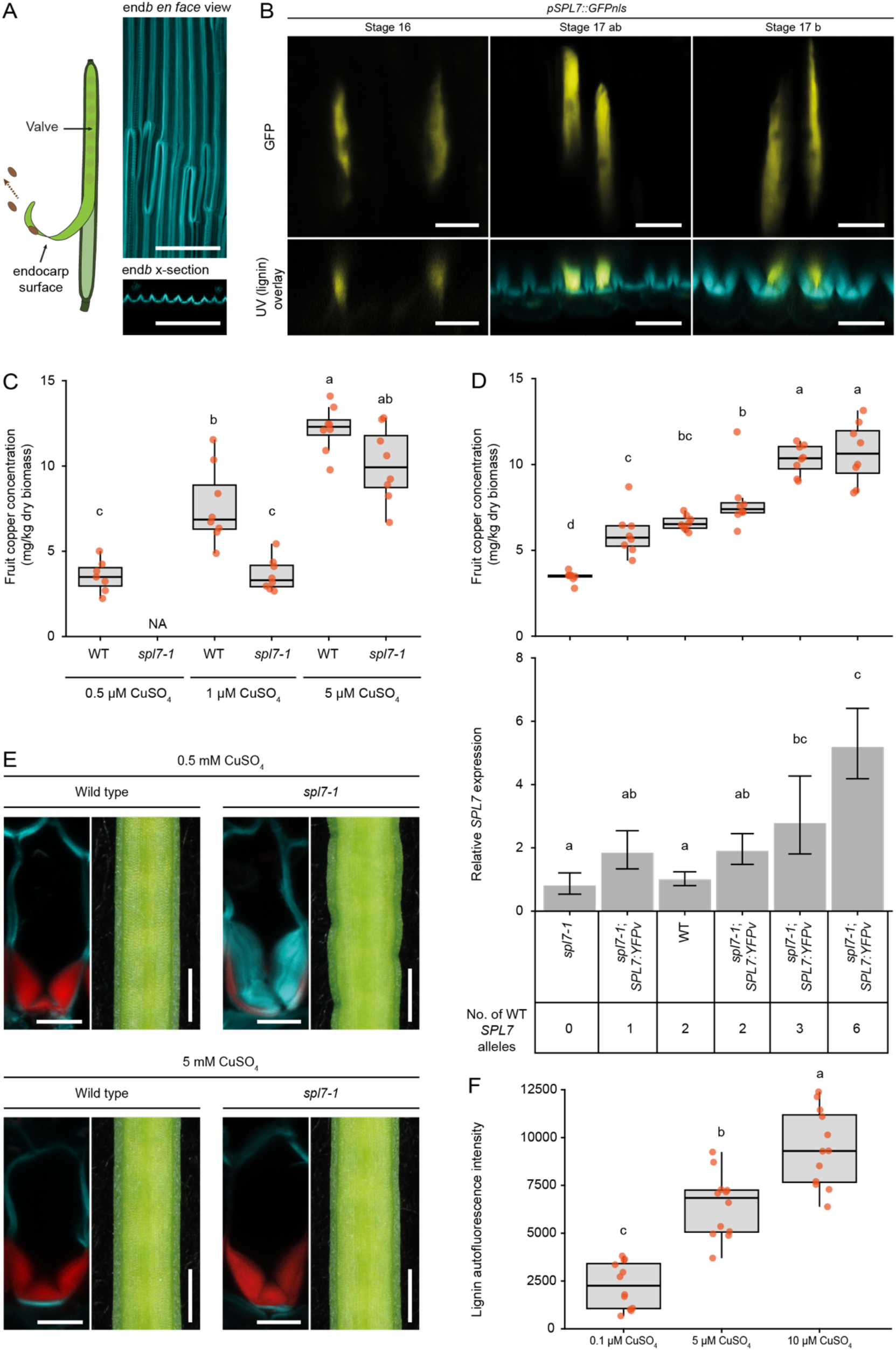
SPL7 is necessary for Cu accumulation in *C. hirsuta* fruit. (*A*) Schematic indicating the endocarp surface of the fruit valve (*left*) and end*b* SCWs imaged *en face* (*top right*) and shown as transverse optical sections (*bottom right*) to indicate the orientation of panels in (*B*). Lignin autofluorescence shown in cyan. (*B*) *pSPL7::GFPnls* expression (yellow) in end*b* cell nuclei of stage 16, 17ab and 17b fruit valves, imaged *en face* (*top*) and shown together with lignin autofluorescence (cyan) in transverse optical sections (*bottom*). (*C*) Boxplot of Cu concentration (mg/Kg dry biomass) in mature fruits of wild-type and *spl7-1* plants grown in an aeroponics system and irrigated with solutions containing either 0.5, 1 or 5 µM CuSO_4_ ; n ≥ 7 biological replicates per condition (red dots) where each replicate contains 5 to 7 pooled fruits from multiple plants; NA (not applicable) indicates the absence of *spl7-1* fruit in this condition. (*D*) Dose-response between *SPL7* gene expression and Cu concentration in mature fruit of *spl7-1*, wild type and *spl7-1; SPL7:YFPv* complementation lines with an increasing number of wild-type *SPL7* alleles. Boxplot of Cu concentration (mg/kg dry biomass) in mature fruit; n = 8 biological replicates per genotype (red dots) where each replicate contains 5 pooled fruits from multiple plants (*top*). *SPL7* gene expression in mature fruit of the same genotypes measured by quantitative RT-PCR, normalized against the housekeeping gene *TIP41* (CARHR242510), and expressed relative to wild type; error indicates SD; n = 3 biological replicates per genotype where each replicate contains two pooled fruit (*bottom*). The number of wild-type *SPL7* alleles present in each genotype is indicated. (*E*) End*b* cells and fruit margins of wild type and *spl7-1* mutant. All plants supplemented with 0.5 mM CuSO_4_ for 4 weeks; 5 mM CuSO_4_ provided as additional supplementation during fruit development. Lignin stained with basic fuchsin (red), cell walls stained with calcofluor white (cyan). (*F*) Boxplot of lignin autofluorescence in end*b* cells of *spl7-1* fruit excised and grown on MS media containing either 0.1, 5 or 10 µM CuSO_4_ ; n = 12 end*b* cell regions per treatment (red dots) from 3 fruits from multiple plants. Plants were grown on soil and supplemented with 0.2-0.5 mM CuSO_4_ to ensure growth and development before fruit were excised. Boxplots show medians (thick black lines), different letters denote statistical significance at *P* < 0.05 using one-way ANOVA and Tukey’s test as post hoc analysis (*D, F*) or Kruskal-Wallis and Fisher’s least significant difference as post hoc analysis (*C*). Confocal micrographs show z-axis sum projections of fruit valves *en face* (*A-B*) or transverse sections (*E*). Scale bars: 100 µm (*A*), 20 µm (*B*), 10 µm (*E, Left*) and 1 mm (*E, Right*).

### SPL7 regulates Cu concentration in fruit

In *A. thaliana, SPL7* is required for plants to acclimate to Cu deprivation (29). To test this in *C. hirsuta*, we grew wild-type and *spl7* plants on soil in low Cu conditions and found that plant size was severely reduced in *spl7*, but unaffected in wild type (Fig. S2). To test whether SPL7 regulates Cu homeostasis in *C. hirsuta* fruit, we grew plants in aeroponic chambers to directly supply bioavailable Cu, and measured Cu concentration in the fruit by ICP-MS. We found that *spl7* fruit accumulated significantly less Cu than wild type under low Cu conditions (1 µM CuSO_4_, Fig. 2C). In contrast to wild type, no fruit were produced by *spl7* plants in 0.5 µM CuSO4, indicating that minimal Cu supplementation is required to restore *spl7* fertility (Fig. 2C) (32, 33). The concentration of Cu in both wild-type and *spl7* fruit increased in response to the concentration of Cu supplied to the roots, over a range of 0.5 – 5 µM CuSO_4_ (Fig. 2C). At high concentrations of Cu, the difference in fruit Cu concentration was no longer significant between genotypes, indicating that sufficient Cu supplementation can bypass the need for *SPL7* (Fig. 2C). Transgenic expression of SPL7 fully restored the Cu concentration in *spl7* fruit to wild-type levels (Fig. 2D, Fig. S2). Using six different *SPL7* genotypes grown in Cu-limiting conditions, we found a dose response between increasing SPL7 activity and increasing Cu levels in the fruit (Fig. 2D). Therefore, *SPL7* is necessary for Cu to accumulate in fruit tissues in low Cu conditions, and is also sufficient to increase Cu above wild-type levels in the fruit.

The thick, asymmetric, end*b* SCW in *C. hirsuta* is built up by sequential deposition of lignified SCW layers (Fig. 1F) (4). In *spl7* fruit, we observed variable amounts of lignin in individual SCW layers, suggesting that SPL7 likely buffers lignification against fluctuations in Cu availability (Fig. 1G). To understand whether different phenotypes of the *spl7* mutant are all attributable to defective Cu homeostasis, we grew plants on soil in high Cu conditions. This rescued the buckling of the fruit margin and the reduced lignification of end*b* cell walls and caused a significant increase in the height and biomass of *spl7* mutants (Fig. 2E, Fig. S2). To directly quantify the effect of Cu concentration in the fruit on lignin deposition in end*b* cells, we grew *spl7* fruit in Cu-containing media. We observed a dose response between increasing Cu concentration and increasing amounts of lignin autofluorescence in end*b* SCWs over a range of 0.1 to 10 µM CuSO_4_ (Fig. 2F). Therefore, the mechanism of localized lignin deposition in end*b* cells is conditional on SPL7 activity and shows a dose dependence on Cu concentration in the fruit.

### Polar localization of LAC4, 11 and 17 precisely predicts end*b* lignification

To identify genes required for end*b* lignification, we took advantage of the less lignified fruit valves of *spl7* mutants for RNA sequencing. Among the ten most significant differentially expressed genes between *spl7* and wild-type fruit valves, are two genes encoding lignin polymerizing enzymes (Fig. 3A, Dataset S1). *LACCASE 11* (*LAC11*, CARHR210090) and *PEROXIDASE 49* (*PER49*, CARHR244760) are both significantly upregulated in *spl7* (Fig. 3A, Fig. S3). On the other hand, phenylalanine ammonia-lyase (PAL, CARHR084820) and cinnamate-4-hydroxylase (C4H, CARHR123940) – genes encoding enzymes that catalyse the first steps of monolignol biosynthesis – are significantly downregulated in *spl7* (Fig. 3A, Fig. S3). To investigate whether reduced monolignol biosynthesis might contribute to the reduced lignification of *spl7* end*b* cells, we grew *spl7* fruit in low Cu media containing monolignols (coniferyl and sinapyl alcohols). The addition of monolignols had no effect on end*b* SCW lignification (Fig. S3), suggesting that the downregulation of *PAL* and *C4H* gene expression in *spl7* fruit valves is not the cause of less lignin in end*b* cells but rather a consequence of feedback mechanisms. The upregulation of *LAC11* and *PER49* gene expression in *spl7* fruit valves may reflect similar feedbacks.

**Figure 3.**
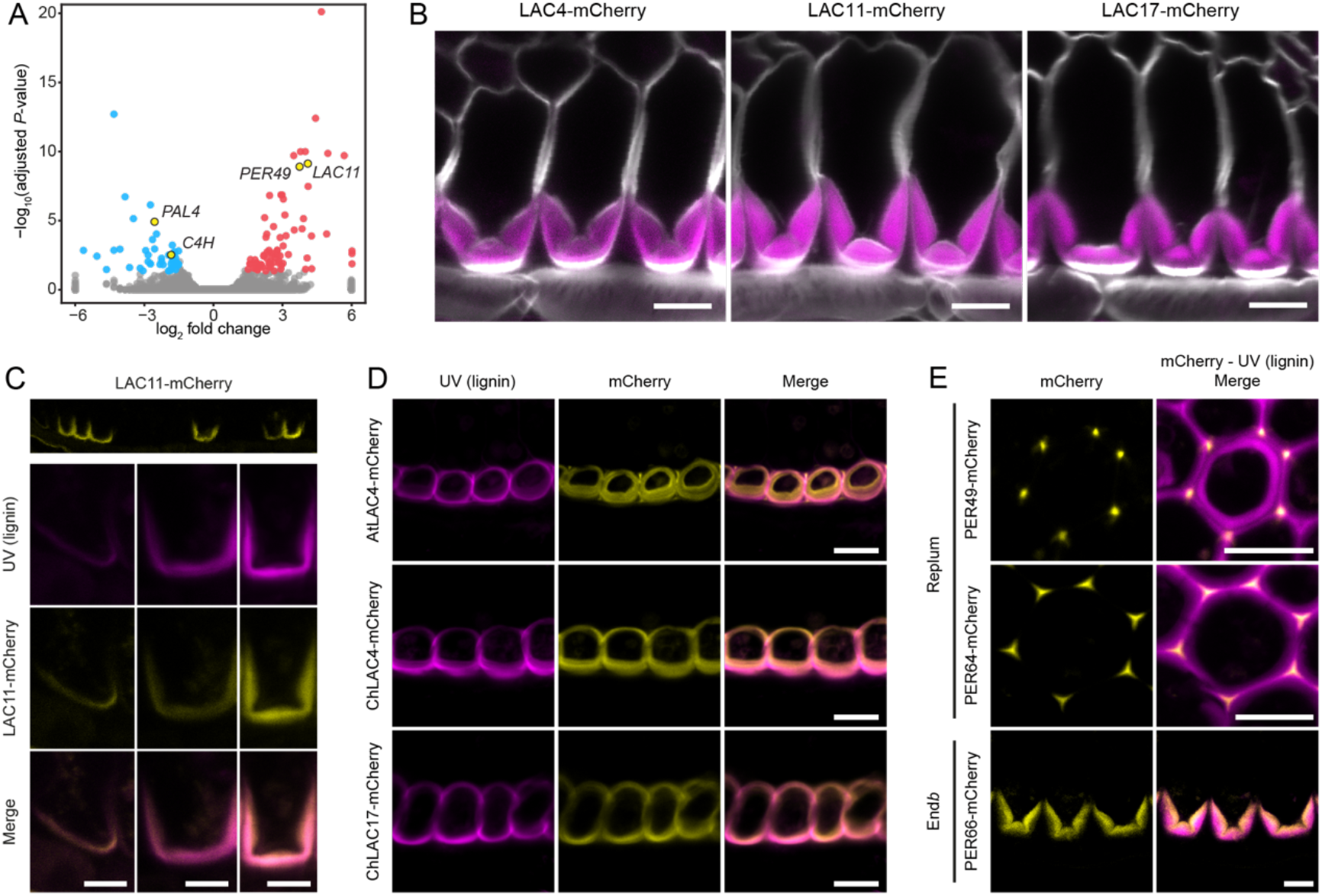
LACCASE4, 11 and 17 co-localize with lignin in end*b* cell walls. (*A*) Volcano plot of differential gene expression between wild-type and *spl7-1* fruit valves with up-regulated genes (log2-fold change > 1.5) shown in red and down-regulated genes (log2-fold change < -1.5) shown in blue; adjusted *P*-value < 0.05. Genes related to lignin biosynthesis indicated in yellow. For visualization purposes, values of log2 fold change <-6 or >6 were set to -6 and 6, respectively. (*B*) *C. hirsuta* LAC4, LAC11 and LAC17 protein fusions (magenta, *pLAC::LAC:mCherry*) localize to *C. hirsuta* end*b* SCWs, cell walls stained with calcofluor white. (*C*) *pLAC11::LAC11:mCherry* expression (yellow) in *C. hirsuta* end*b* SCWs as they start to lignify (*top*) and close-up views of individual end*b* SCWs (*bottom*) shown together with lignin autofluorescence (magenta). (*D*) *A. thaliana* LAC4 and *C. hirsuta* LAC4 and LAC17 protein fusions (yellow, *pLAC::LAC:mCherry*) co-localize with non-polar lignin autofluorescence (magenta) in *A. thaliana* end*b* SCWs. (*E*) *C. hirsuta* PER49, PER64 and PER66 protein fusions (yellow, *pPER::PER:mCherry*) co-localize with lignin autofluorescence (magenta) in replum cells (PER49, PER64) or end*b* cells (PER66) in *C. hirsuta* fruit. Confocal micrographs show z-axis sum projections of transverse fruit sections (*B-E*). Scale bars: 10 µm (*B, D, E*), 5 µm (*C*)

Interestingly, laccases are multi-Cu glycoproteins and thus good candidates to control Cu-dependent lignin deposition in end*b* cells. We detected transcripts for 6 out of 15 *LAC* gene family members in *C. hirsuta* fruit valves (mean of wild type and *spl7-1* normalized read counts > 5), with *LAC4* and *LAC17* showing the highest expression (Fig. S3). Type III peroxidases are heme-containing glycoproteins, encoded by a large gene family with 62 members annotated in the current *C. hirsuta* genome assembly (34). We detected transcripts for 9 out of 62 *PER* gene family members *C. hirsuta* fruit valves (mean of wild type and *spl7-1* normalized read counts > 5), including *PER64* which is required for Casparian strip lignification in *A. thaliana* (12, 16), with *PER42* and *PER66* showing the highest expression in wild type (Fig. S3). We selected *LAC4, LAC11, LAC17, PER49, PER64* and *PER66* for further study and generated reporters to visualize where and when these genes and their protein products are expressed in *C. hirsuta* fruit.

Similar to *SPL7*, we found expression of *LAC4, LAC11* and *LAC17* (*pLAC::GFPnls*) in end*b* cells in the fruit valve (Fig. S3). Given the asymmetric deposition of lignin within *C. hirsuta* end*b* SCWs, we predicted that enzymes required for lignin polymerization should pre-pattern this asymmetry. Fluorescent protein fusions of LAC4, LAC11 and LAC17 (*pLAC::LAC:mCherry*) precisely co-localized with asymmetric lignin deposits in end*b* SCWs (Fig. 3B). LAC11 localized preferentially to lignified cell walls in the valve of the fruit, while LAC4 and LAC17 also localized to lignified cell walls in the replum (Fig. S3). LAC11 first accumulated asymmetrically in a thin layer of the end*b* SCW, forming a U-shape in cross-section (Fig. 3C). This asymmetric pattern co-localized perfectly in time and space with lignin (Fig. 3C), indicating that the required monolignols are present as substrates in the apoplast for lignin polymerization. LAC11 continued to accumulate throughout the U-shaped SCW as it rapidly thickened and started to form characteristic hinges at the base of the U (Fig. 3C). We next investigated whether these *C. hirsuta* laccases maintained their polar localization when transferred into *A. thaliana*. In the non-explosive fruit of *A. thaliana, pAtLAC4::AtLAC4:mCherry* co-localized with the symmetrically lignified end*b* SCWs (Fig. 3D). Similarly, we observed a symmetric localization of *C. hirsuta* LAC4 and 17 fusion proteins in *A. thaliana* end*b* cell walls, but we could not detect *C. hirsuta* LAC11 (Fig. 3D, Fig. S3). These results suggest that the asymmetric localization of *C. hirsuta* LAC4 and 17 is not determined in *cis* to these genes, but depends on additional polarity determinants that are present in *C. hirsuta*, but not *A. thaliana*, end*b* cells.

To understand whether peroxidases show a similar pattern to laccases, we localized mCherry C-terminal fusions of PER49, PER66 and PER64 (*pPER::PER:mCherry*) in *C. hirsuta* fruit (Fig. 3E). We could not detect PER49 or PER64 in end*b* SCWs in the valve, but rather in lignified cell walls in the replum. PER49 and PER64 localized to the middle lamella at cell corners and did not accumulate throughout the lignified SCWs (Fig. 3E). However, PER66 localized precisely to end*b* SCWs in the valve of the fruit, similar to LAC11 (Fig. 3E, Fig. S3). Therefore, all three laccases, but only one of three peroxidases that we examined, localized to end*b* SCWs.

### *LACCASE 4* and *17* are required for end*b* lignification

To directly assess whether *LAC4, 11* and *17* are needed for end*b* lignification, we generated loss-of-function alleles for all three genes using a CRISPR/Cas9 multiplex guideRNA strategy. Premature stop codons in these alleles result in truncated proteins that lack aa residues required to coordinate the Cu ions that are essential for laccase activity (Fig. S4). These triple mutants, therefore, knock out *LAC4, 11* and *17* function and show severe growth arrest, which can be partially rescued in culture to produce stems with reduced lignification and collapsed xylem vessels (Fig. 4A, Fig. S5). Dexamethasone induction of a *pLAC11::LhGR>>LAC11:mCherry* construct (*pLAC11::GR-LhG4/pOp6::LAC11:mCherry*) was sufficient to rescue the growth arrest of triple *lac4 11 17* mutants, demonstrating that this phenotype is LAC-dependent (Fig. 4A).

**Figure 4.**
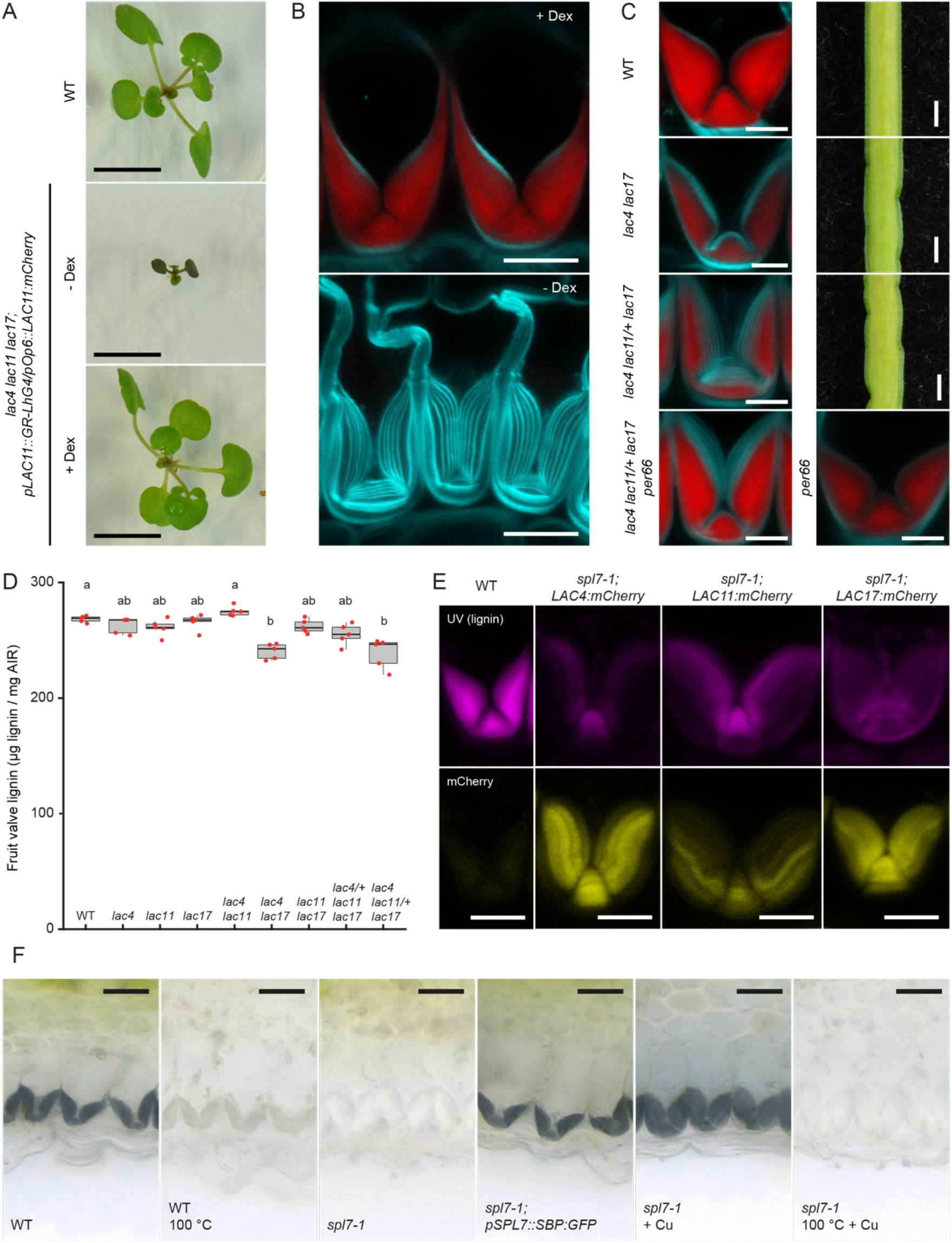
Cu-dependent laccase activity is necessary for end*b* lignification in *C. hirsuta* fruit. (*A-B*) Triple *lac4 11 17* mutants expressing *pLAC11::LhGR>>LAC11:mCherry*. Two-week old seedlings grown on ½ MS plates without (-Dex) or with 10 µM dexamethasone (+ Dex), compared to wild type (*A*). End*b* cells of plants treated with 100 µM dexamethasone daily until fruit matured (+ Dex) or only until plants started to bolt, such that fruit developed in the absence of dexamethasone (- Dex) (*B*). Lignin, stained red with basic fuchsin, cell walls stained cyan with calcofluor white. (*C*) End*b* cells and close-up view of fruit margin in wild-type, *lac4 17* and *lac4 11/+ 17*. End*b* cells in *lac4 11/+ 17 per66* and *per66* fruit. Lignin, stained red with basic fuchsin, cell walls stained cyan with calcofluor white. (*D*) Boxplot of lignin concentration shown as µg of acetyl bromide soluble lignin per mg of alcohol-insoluble residue (AIR) in mature fruit valves of wild type and allelic combinations of *lac4, lac11* and *lac17* mutants. Plot shows median (thick black line), n = 5 biological replicates per genotype (red dots) where each replicate contains 16 valves from two plants, different letters denote statistical significance at *P* < 0.05 using Kruskal-Wallis and Dunn’s test as post hoc analysis. (*E*) LAC4, LAC11 and LAC17 protein fusions (yellow, *pLAC::LAC:mCherry*) accumulate in *spl7-1* end*b* SCWs, which have less lignin (magenta, autofluorescence) than wild type. (*F*) Laccase activity (indicated by black precipitate produced by oxidation of 4-hydroxyindole substrate) in fruit valves of wild type (± enzyme inactivation by heat), *spl7-1, spl7-1; pSPL7::SBP:GFP, spl7-1* incubated with 5 mM CuSO_4_ (± enzyme inactivation by heat). Confocal micrographs show z-axis sum projections of transverse fruit sections (*B-C, E*). Scale bars: 10 mm (*A*), 10 µm (*B, E*), 5 µm (*C*), 1 mm (fruit, *C*), 20 µm (*F*).

To observe end*b* SCW lignification in *lac4 11 17* triple mutants, we took advantage of the rescue conferred by *pLAC11::LhGR>>LAC11:mCherry* to grow triple mutants through to flowering. By removing dexamethasone induction at this point, we recovered *lac4 11 17* fruit that no longer expressed LAC11:mCherry. End*b* SCWs in these fruits failed to lignify, whereas continued dexamethasone induction produced fully lignified end*b* SCWs (Fig. 4B). Therefore, LAC4, 11 and 17 are redundantly required for localized lignin deposition in end*b* cells of *C. hirsuta*. We used various allelic combinations of *lac4, 11* and *17* to assess their relative contribution to end*b* lignification. We found significantly less lignin in the fruit valves of *lac4 17* double mutants and *lac4 11/+ 17* plants that contained five mutant alleles (Fig. 4D). We observed lignin in only a portion of the end*b* SCW in these genotypes (Fig. 4C). Newly deposited SCW layers lacked lignin in a similar way to Cu-deprived fruit of *spl7* mutants (Fig. 1C, G). Therefore, *LAC4* and *17* contribute redundantly to end*b* lignification and *LAC11* is required with both genes to fully lignify the end*b* SCW. We found additional similarities between the fruit phenotypes of *spl7* mutants and *lac4 17* and *lac4 11/+ 17* mutants. For example, buckling occurred along the edges of mature fruit in both *lac* genotypes (Fig. 4C), and these fruit valves had significantly elevated ratios of syringyl to guaiacyl (S/G) lignin monomers, which was also found in *spl7* alleles, and indicative of laccase deficiency (Fig. S5) (17). Therefore, the Cu-dependence of end*b* lignification in *spl7* fruit, is likely to reflect the contribution of multi-copper laccases to polymerizing lignin in this cell type.

PER66 showed a similar, polar localization to LAC4, 11 and 17 in the SCW of *C. hirsuta* end*b* cells (Fig. 3E). To investigate any additional contribution of this peroxidase to end*b* SCW lignification, we used CRISPR/Cas9 to generate a *per66* loss-of-function allele in a segregating *lac4 lac11/+ lac17* background. A premature stop codon in *per66-1* resulted in a truncated 121 aa protein (Fig. S4). The defects in end*b* SCW lignification observed in *lac4 lac11/+ lac17* were not enhanced by *per66*, and *per66* mutants did not differ from wild type (Fig.4C). Moreover, end*b* lignification was unaffected by addition of the peroxidase inhibitor salicylhydroxamic acid (SHAM) (12, 35) when we grew *spl7* fruit in Cu-containing media (Fig. S3). Therefore, the highly localized patterns of LAC4, 11 and 17, but not PER66, are required for end*b* SCW lignification. This is reminiscent of Casparian strip formation, where it is laccases that are replaceable for lignification, while peroxidases are absolutely required (16). In both cases, genetic evidence indicates that despite co-localization, only one class of oxidative enzymes is required for local lignin deposition.

### SPL7-dependent Cu homeostasis is required for laccase activity in end*b* SCWs

Our findings suggest that SPL7 is needed to ensure sufficient Cu in the fruit for the activity of LAC4, 11 and 17 in end*b* cell walls to form lignin. To test this hypothesis, we first verified that LAC4, 11 and 17 protein fusions accumulate in the end*b* cell walls of *spl7* fruit (Fig. 4E). We also verified that the laccase-specific oxidation of 4-hydroxyindole was suitable to detect laccase activity in glycoprotein extracts of wild-type fruit (Fig. S5). Using this assay *in situ*, we detected high levels of laccase activity localized to end*b* SCWs in wild-type fruit (Fig. 4F). Heat inactivation showed that this activity is enzyme-dependent (Fig. 4F). We found no laccase activity in the end*b* SCWs of *spl7* fruit grown in low Cu conditions (Fig. 4F). The absence of laccase activity matched the reduced lignin in these SCWs (Fig. 1C). Laccase activity was restored in *spl7* end*b* SCWs by complementation with a *pSPL7::SBP:GFP* transgene (Fig. 4F). Strikingly, enzyme-dependent laccase activity was restored in *spl7* end*b* SCWs by direct application of CuSO_4_ to *spl7* fruit tissue (Fig. 4F). This result indicates that although laccases are present in *spl7* end*b* SCWs, they require Cu supplementation for enzymatic activity. Therefore, localized lignin deposition in end*b* SCWs requires three laccases (LAC4, 11 and 17), which depend on SPL7 to provide sufficient Cu for their activity.

## Discussion

Explosive seed dispersal in *C. hirsuta* depends on the precise sub-cellular deposition of lignin in end*b* cells of the fruit valves. Here, we identified four lignin polymerizing enzymes – PER66, LAC4, 11 and 17 – that co-localize with the asymmetric pattern of lignin in end*b* SCWs. We used conditional gene expression to bypass the growth arrest of *lac4 11 17* triple mutants and show that LAC4, 11 and 17 are required to lignify the distinctive end*b* SCW. The requirement for multi-Cu laccases rather than peroxidases to polymerize lignin in this cell type, explains why SPL7, a key regulator of Cu homeostasis, is essential for robust end*b* SCW lignification.

The polar deposition and hinged pattern of lignin in end*b* cells of explosive fruit is an evolutionary novelty of *Cardamine*, associated with the appearance of explosive seed dispersal in this genus of the Brassicaceae family (4). We found that *C. hirsuta* LAC4 and 17 adopted a non-polar localization when expressed under their native promoters in *A. thaliana*, matching the non-polar lignification of *A. thaliana* endb SCWs. These findings show conservation between species in *LAC* gene expression, but not protein localization. Therefore, the polarity and pattern of *C. hirsuta* LAC4, 11 and 17 end*b* localization, is likely to be determined by *trans* factors that are specific to *C. hirsuta*. Identifying such factors will be an important follow-up to this study.

Apoplastic laccases are synthesized in the endomembrane system and are secreted by exocytosis (36). Hence, targeting laccase-loaded vesicles to specific regions of the plasma membrane may be important to pattern their localization in the cell wall. Laccases are also immobile in the dense SCW matrix and their localization may be fixed by anchoring to specific SCW components (11). LAC11 localization in *C. hirsuta* end*b* SCWs is coincident with the initial, asymmetric deposition of lignin throughout the SCW. In contrast to this, *C. hirsuta* LAC4, 17 and PER66 accumulate to higher levels in distinct layers of the lignified end*b* SCW. This may reflect spatial regulation within the SCW or temporal regulation during the sequential deposition of SCW layers in *C. hirsuta* end*b* cells (4). Interestingly, dexamethasone induction of *pLAC11::LhGR>>LAC11:mCherry* can overcome the restriction of LAC11 to the polar SCW domain and cause ectopic lignification throughout the end*b* cell wall (Fig. S5). This suggests that levels of *LAC11* expression in end*b* cells can influence protein localization, and further suggests that monolignols are available for lignin polymerization throughout the end*b* cell wall. It is an open question whether the hinged pattern of lignin deposition in *C. hirsuta* end*b* cells is also regulated by the same factors that determine polarity, or by an independent mechanism.

The reduced range of seed dispersal in *spl7* mutants suggests that the material properties of lignin in the end*b* cell wall influence explosive dispersal. The tension that generates elastic energy for explosion is produced by differential contraction of valve tissues in *C. hirsuta* fruit (4). This puts the end*b* layer under compression and the exocarp layer under tension, and results in the storage of potential elastic energy. Material properties of the end*b* SCW will determine its compressive strength. We observed that less lignified end*b* SCWs in *spl7, lac4 17* and *lac4 11/+ 17* genotypes, resulted in buckling of the fruit valve along its margin under load. Consequently, the amount of stored potential elastic energy released in this buckling would no longer be available for explosive valve coiling. This offers one explanation for the reduction in seed dispersal range in *spl7* mutants. Future experiments will be useful to distinguish between alternative hypotheses.

Homeostatic regulation of Cu is a conserved function of SPL7 between *C. hirsuta* and *A. thaliana*. In *A. thaliana, LAC4* and *17*, but not *LAC11*, are targeted by SPL7-activated Cu miRNAs for post transcriptional degradation in response to Cu deprivation (28). Although this regulation may also be conserved in *C. hirsuta*, our findings show that it is laccase activity, rather than the abundance of LAC4 and 17, that determines Cu-dependent end*b* lignification in *spl7*. Future work to dissect the SPL7 transcriptional response to Cu deprivation in *C. hirsuta* fruit will help to identify the precise mechanisms through which Cu is made available for LAC4, 11 and 17 activity in end*b* SCWs via the SPL7 pathway. The Cu-dependence of laccase activity means that lignification of individual layers of end*b* SCWs in *C. hirsuta spl7* fruit provides a cell-level read-out of Cu availability (Fig. 1F). This read-out gives some insight into the large variation in Cu availability that cells experience during growth and development, in the absence of SPL7. This variation in Cu availability in different plant tissues, throughout development, and between different growing conditions, is buffered by SPL7 activity. The functional conservation of SPL7 homologs from green algae to flowering plants is indicative of how essential this regulation of Cu homeostasis is for green plant life (22, 26, 29).

In summary, we have identified a module that links mineral nutrition with seed dispersal. Localized lignin deposition is critical for the mechanism of explosive seed dispersal in *C. hirsuta*. Cu-requiring laccases regulate this lignification, making explosive dispersal dependent on the homeostatic control of Cu by SPL7. In this way, a SPL7/LAC4/11/17 module integrates mineral nutrition with polar lignin deposition to facilitate dispersal.

## Materials and Methods

### Plant material

*Cardamine hirsuta* (Ox), herbarium specimen voucher Hay 1 (OXF) (3), and *Arabidopsis thaliana* (Col-0) were used as wild type. The *lig1* (*spl7-1*) allele was identified in a previous mutant screen in *C. hirsuta* (4), and the causal mutation identified by mapping-by-sequencing. The following mutant alleles were generated in *C. hirsuta* using CRISPR/Cas9: *spl7-2, per66-1, lac4-1, lac4-2, lac11-1* and *lac17-1* (see *SI Appendix* for more details). *pAtLAC4::AtLAC4:mCherry; lac4 lac17* was previously described (8). The following transgenic lines were generated in *C. hirsuta* for this study (constructs described in *SI Appendix*): *pSPL7::GFP-NLS, pSPL7::mCherry:SPL7, pSPL7::ΔSPL7(SBP):GFP, pLAC11::LhGR::pOp6::LAC11:mCherry, pLAC4::GFP-NLS, pLAC11::GFP-NLS, pLAC17::GFP-NLS, pLAC4::LAC4:mCherry, pLAC11::LAC11:mCherry, pLAC17::LAC17:mCherry, pPER49::PER49:mCherry, pPER64::PER64:mCherry, pPER66::PER66:mCherry*. Plants were grown on soil, *in vitro* and using aeroponics, both with and without CuSO_4_ supplementation. Dexamethasone was supplied in solid media and spray solution. Fruits were grown *in vitro* and treated with CuSO_4_, peroxidase inhibitor Salicylhydroxamic acid, and the monolignols coniferyl alcohol and sinapyl alcohol. More details are described in *SI Appendix*.

### Seed dispersal

Seed dispersal distance was measured in concentric rings around wild-type and *spl7-1* plants as previously described (4) with slight modifications (see *SI Appendix*).

### Microscopy

A Leica TCS SP8 was used for Confocal Laser Scanning Microscopy (CLSM) with the following excitation and emission parameters (nm): lignin autofluorescence ex: 405, em: 440-510; calcofluor ex: 405, em: 425-475; GFP ex: 488, em: 500-550 or 492-540; basic fuchsin ex: 561, em: 600-650; mCherry ex: 594, em: 600-640; Chlorophyll ex: 488, em: 650-730. Epifluorescence and brightfield microscopy were performed using a Zeiss Axio Imager M2 microscope. Cryo-fracture scanning electron microscopy was performed using an Emitech K1250x cryo unit and a Zeiss Supra 40VP microscope. CLSM images were processed, and lignin autofluorescence intensity quantified, using the Fiji package of ImageJ (https://fiji.sc). More details are described in *SI Appendix*.

### Histochemistry

The ClearSee protocol (37) was used to visualize fluorescent proteins and cell wall stains (basic fuchsin and calcofluor white) by CLSM. Phloroglucinol staining of lignin was performed as described (4). Fresh fruit sections were treated with 4-hydroxyindole to visualize laccase activity; pre-treatments included 100°C or CuSO_4_.

### Cu quantification by ICP-MS

Cu content of fruits was measured using an Agilent 7700 ICP-MS and expressed as mg/Kg dry biomass. More details are described in *SI Appendix*.

### Lignin quantification and monomer analysis

Acetylbromide soluble lignin and lignin composition via thioacidolysis was determined as previously described (38). More details are described in *SI Appendix*.

### RNAseq

RNA was extracted from three biological replicates of wild-type and *spl7-1* stage 17 fruit valves using the Spectrum Total RNA kit, and cDNA was synthesized using SuperScript III Reverse Transcriptase. Libraries (100 bp) were prepared and sequenced using the HiSeq2500 Illumina platform at the MPIPZ Genome Centre. Paired-end reads were quality-checked, aligned to the *C. hirsuta* reference genome and quantified (34). Differential gene expression was analyzed using DESeq from Bioconductor (39). More details are described in *SI Appendix*.

### qRT-PCR

RNA was extracted from three biological replicates of stage 17 fruit per genotype and used for cDNA synthesis as described above. qPCR was performed on a QuantStudio5 thermocycler using SYBR Green Supermix. The housekeeping gene *TIP41* (CARHR242510) was used as reference and relative expression of *SPL7* was calculated using the 2^-ΔΔCt^ method. Primers used are listed in *SI Appendix*, Table S1.

### Statistical Analyses

Statistical analyses were done with R Statistical Software (40). More details are described in *SI Appendix*.

## Data Availability

Short sequence reads are available at European Nucleotide Archive in bioproject PRJEB50935. All other data are included in the article and/or SI Appendix.

## Acknowledgments and funding sources

We thank P. Huijser, M. Tsiantis and A. Emonet for comments, K. Lufen for lignin analyses, P. Sarchet for conducting the mutant screen, L. Samuels, A. Maizel and C. Kamei for sharing materials, X. Gan for bioinfomatic services, A. Stamatakis for greenhouse support, R. Franzen for SEM and W. Faigl for laccase purification. This work was supported by an IMPRS studentship to M.P-A., the Deutsche Forschungsgemeinschaft (DFG) under Germany’s Excellence Strategy— EXC 2048/1—Project ID: 390686111 to M.P. and DFG FOR2581 Plant Morphodynamics grant to A.H. Portions of the paper were developed from the thesis of M.P-A.

## Notes

### Competing Interest Statement

The authors have declared no competing interest.

