## Supplementary Information for "Explosive seed dispersal depends on SPL7 to ensure sufficient copper for localized lignin deposition via laccases"

Current address: [4] University of Cologne, Weyertal 115c, 50931 Cologne, Germany; [5] Keygene N.V., 6700 AE Wageningen, The Netherlands.

**This PDF file includes:**

SI Text
Tables S1 to S2
Figures S1 to S5
SI References

### Supplementary Information Text

### Supplementary materials and methods

#### Plasmid construction

*pSPL7::GFP-NLS*, *pSPL7::mCherry:SPL7*, *pSPL7::ΔSPL7(SBP):GFP*, *pLAC4::GFP-NLS*, *pLAC11::GFP-NLS*, *pLAC17::GFP-NLS*, *pLAC4::LAC4:mCherry*, *pLAC11::LAC11:mCherry*, *pLAC17::LAC17:mCherry*, *pPER49::PER49:mCherry*, *pPER64::PER64:mCherry* and *pPER66::PER66:mCherry* were generated by GreenGate cloning, which relies on the *Bsa*I enzyme, therefore, any *Bsa*I sites were mutagenized as described (1). The following promoters were PCR-amplified (base pairs [bp] before ATG): *pSPL7* (2,374), *pLAC4* (3,696), *pLAC11* (3,252), *pLAC17* (2,885), *pPER49* (958 [whole intergenic region]), *pPER64* (2,888) and *pPER66* (3,292). Full gene coding sequences were amplified to generate translational fusions. *pSPL7::ΔSPL7(SBP):GFP* was generated by amplifying only the first 759 bp of *SPL7* coding sequence. Entry vector combinations were cloned into binary vectors pGGZ003 or pGGZwf01, a GreenGate-compatible pPZP200-based vector (2) generated in this study. *pLAC11::GR-LhG4/pOp6::LAC11:mCherry* was generated as a multiple expression GreenGate cassette using the additional entry vectors pOp6 and GR-LhG4 (3). Entry vectors were subsequently cloned in pGGZwf01 binary vector. *pSPL7::SPL7:vYFP* was generated by MultiSite gateway (Thermo Fisher Scientific) using plasmids generated in (4). A 2,374 bp promoter fragment was recombined in the entry vector pGEMt-easy1R4. The full SPL7 coding sequence was recombined in the entry vector pGEMt-easy221. These two entry vectors were recombined with the entry vector YFP-nosT pGEMt-easy2R3 into the pGREENII125 binary vector.

#### Plant material and growth conditions

*Cardamine hirsuta* (Ox), herbarium specimen voucher Hay 1 (OXF) (5), and *Arabidopsis thaliana* (Col-0) were used throughout this study unless otherwise specified. Transgenic *C. hirsuta* and *A. thaliana* plants were generated by the floral dip method (6) using *Agrobacterium tumefaciens* strains GV3101 or C58. The *A. thaliana* transgenic line *pAtLAC4::AtLAC4:mCherry; lac4 lac17* was a kind gift from L. Samuels (7). The *spl7-2*, *per66-1*, *lac4-1, lac4-2, lac11-1* and *lac17-1* alleles were generated in *C. hirsuta* by CRISPR/Cas9 gene editing (8). The *lig1* (*spl7-1*) allele was identified in a previous EMS screen in *C. hirsuta* Ox (9). A homozygous *lig1* mutant was crossed to the *C. hirsuta* Nz accession (6) to generate a F2 mapping population. DNA was extracted from tissue of 48 *lig1* plants from the F2 mapping population and sequenced at the MPIPZ genome centre on an Illumina HiSeq 2000 platform. Paired-end reads were mapped to the Ox reference genome and single nucleotide polymorphisms

(SNPs) were called against the reference sequence. To confirm the mapping-by-sequencing results and to narrow down the causal interval, seven markers (Table S1) were analysed in individual *lig1* recombinants to fine-map the *lig1* mutation between positions 17676369 bp and 17882576 bp on chromosome 6. In this interval, two nonsynonymous EMS-derived mutations were identified. SNPs derived from the Nz accession were excluded by comparison to genome resequencing data.

Plants were grown at the MPIPZ, Cologne, Germany. Plants grown on soil were cultivated in greenhouse or controlled environment chambers in long-day conditions (LD; days: 20 °C, 16 h; nights: 18 °C, 8 h). Short-day conditions (SD; days: 20 °C, 8 h; nights: 18 °C, 16 h) were used as specified. *C. hirsuta* and *A. thaliana* seeds were stratified on soil at 4°C in the dark for 7 days and 3 days respectively. For copper supplementation on soil, 10 ml $CuSO_4$ solution was applied to the soil surface once per week, for 4 weeks from time of seed sowing, unless otherwise stated. For experiments in the aeroponic system, plants were grown in small permeable pots filled with inert perlite, and inserted through the lid of enclosed tanks (EZClone). Nutrient solution was sprayed inside the box four times a day for 15 min using an Aquarius Universal 180 pump (OASE) and 12 spray nozzles (HortiPots). Irrigation solutions contained all nutrients typically used in normal greenhouse conditions and variable concentrations of $CuSO_4$ as specified. pH was kept in a range of 5.5 – 6.0. Seeds were surface-sterilized and plated on ½ Murashige and Skoog (MS) media supplemented with 0.5 µM $CuSO_4$ and then transferred at seedling stage to the pots in the aeroponics system. For *in vitro* cultivation, seeds were surface-sterilized and plated on ½ MS media. Seeds were stratified at 4°C in the dark for 7 days and then grown in LD conditions. For dexamethasone-induction of *lac4 lac11 lac17 pLAC11::LhGR>>LAC11:mCherry* on plates, 10 mM stock solution was added to the media for a final concentration of 10 µM dexamethasone (Sigma-Aldrich; CAS-No. 50-02-2). For extended dexamethasone induction, seeds were germinated on ½ MS media with 10 µM dexamethasone and then seedlings were transplanted to soil. Plants were grown in SD conditions and switched to LD conditions after 4 weeks (specified in figure legend when directly grown in LD conditions). Aerial parts of the plant were sprayed daily with a solution of 100 µM dexamethasone and 0.01 % Triton X-100. To assess the fruit phenotype of *lac4 lac11 lac17* triple mutants, dexamethasone application was stopped when plants started to bolt, such that fruit developed in the absence of *pLAC11::LhGR>>LAC11:mCherry* induction. To assess the rescued fruit phenotype of *pLAC11::LhGR>>LAC11:mCherry; lac4 lac11 lac17* plants, dexamethasone application was prolonged until fruit maturation.

For cultivation of fruit *in vitro*, stage 16 fruit (before SCWs are deposited in end*b* cells) were excised from plants, inserted upright in ½ MS media containing either 0.1, 5 or 10 µM $CuSO_4$ and grown until stage 17 ab. For inhibition of peroxidase activity, ½ MS media contained 150 µM Salicylhydroxamic acid (SHAM; Sigma-Aldrich; CAS-No. 89-73-6). To test for restoration of lignification with exogenous application of monolignols, 100 µM coniferyl alcohol (Sigma-Aldrich; CAS-No. 458-35-5) and 100 µM sinapyl alcohol (Sigma-Aldrich; CAS-No. 537-33-7) were added together to ½ MS media. The solvent DMSO was added in control treatments.

**CRISPR/Cas9 mutagenesis**

For CRISPR/Cas9 directed mutagenesis of *C. hirsuta SPL7*, *LAC4*, *LAC11*, *LAC17* and *PER66*, a similar strategy as in (8) was followed with several modifications. All possible positions for SpCas9 (10) compatible sgRNAs were identified and evaluated using ChopChopv2 (11) or CCTop (12). Two sgRNAs per gene were selected based on location, high efficiency and minimum off-target prediction (Table S2). MultiSite Gateway cloning was used to generate plasmids for CRISPR/Cas9 mutagenesis. One entry vector contained the SpCas9 sequence driven by an egg-cell specific promoter (EC1.2en-EC1.1p) (13). The second entry vector contained the sgRNAs. The two entry vectors were cloned into Gateway-compatible pPZP200-based binary vectors (2) generated in this study, either containing the fluorescence-accumulating seed technology (FAST) red selection marker (pPZP200-FAST-RFP) or the Basta selection marker (pPZP200-BASTA). To target *SPL7*, the two SpCas9 compatible sgRNAs were driven by the *A. thaliana* U6 RNA pol III promoter (8) and the pPZP200-BASTA binary vector was used for plant transformation. A multiplex sgRNA cloning strategy was followed to simultaneously target *LAC4*, *LAC11* and *LAC17*. The six sgRNAs were driven by a single ubiquitin promoter from *Petroselinum crispum* (parsley, pPcUbi) (10) and the pPZP200-RFP-FAST binary vector was used for plant transformation. To target *PER66*, the two SpCas9 compatible sgRNAs were driven by the *A. thaliana* U6 RNA pol III promoter (8). The pPZP200-FAST-RFP binary vector was used to transform wild-type plants and also *lac4 lac11/+ lac17* plants to directly obtain higher order mutants.

**Seed dispersal experiment**

Wild-type and *lig1* (*spl7-1*) plants were grown on soil for 7 weeks, supplemented weekly with 0.2 mM $CuSO_4$ (low Cu conditions). When siliques started to develop, 4 wild-type and 4 *lig1* plants were staked upright and positioned on large, empty tables to measure distance of seed dispersal. At this point, plant height was measured up to the first fruit on the main stem. On average, wild-type plants were 6 cm taller than *lig1* plants, therefore, pots containing *lig1* plants were elevated by 6 cm. This adjustment eliminated the effect of plant height on dispersal distance. Pots were positioned at the centre of concentric rings drawn on a large sheet of plastic at increasing distances (every 25 cm until 200 cm). Additionally, the stem of each plant was passed through a hole in a petri dish placed at the base of the plant to collect seeds falling directly below the plant. Plants were left to disperse their seeds for 6 weeks, then seeds were collected separately from each concentric ring and counted. The tables that were used for the experiment had a width of only 1 m, therefore, for distances further than 1 m, the entire area of the circular ring could not be sampled. For these distance rings, the percentage of area sampled with respect to the total area of the ring was calculated, and the number of seeds was corrected proportional to the area sampled.

**Microscopy**

To generate transverse sections, fruits or stems were embedded in 1.5 ml tubes containing 5-10 % low melting agarose (Hi-Pure Low agarose; Biogene Ltd). 100-150 µm sections were cut with a Leica Vibratome VT1000 S. A Leica TCS SP8 was used for Confocal Laser Scanning Microscopy (CLSM) with objectives HCX PL APO lambda blue (63x/1.20 water) and HC PL FLUOTAR (10x/0.30 dry) and with the following excitation (ex) and emission (em) parameters (wavelength in nm): lignin autofluorescence ex: 405, em: 440-510; calcofluor ex: 405, em: 425-475; GFP ex: 488, em: 500-550 or 492-540; basic fuchsin ex: 561, em: 600-650; mCherry ex: 594, em: 600-640; Chlorophyll ex: 488, em: 650-730. Confocal images were processed using the Fiji package of ImageJ (https://fiji.sc). To quantify lignin autofluorescence intensity, confocal z-stacks were recorded with 12 bit depth avoiding any overexposure. Settings were adjusted for a wild-type sample first and were kept constant for the rest of the experiment. Z-stacks with a similar number of slices were projected using the method "sum slices" and then a threshold was applied to remove non-specific signal outside the region of interest. The value of mean intensity was recorded for the region of interest in each sample. Epifluorescence and brightfield microscopy were performed using a Zeiss Axio Imager M2 microscope with dry objectives EC Plan-Neofluar 40x/0.75 and EC Plan-Neofluar 10x/0.30. Images were acquired either as single-plane images or as z-stacks from which "Extended depth of field" images were generated using Zeiss imaging software ZEN 2. For epifluorescence imaging, filter set 20 HE (excitation BP 546/12 and emission BP 607/80) was used. For scanning electron microscopy, fresh fruit were flash-frozen with liquid nitrogen and subjected to cryo-fracture, sublimation and gold-coating in an Emitech K1250x cryo unit. Images were taken using a Zeiss Supra 40VP scanning electron microscope operating at 3 kV.

**Histological techniques**

To visualize fluorescent proteins and cell wall components using CLSM, the ClearSee protocol was used (14, 15). Lignin was stained with basic fuchsin (0.2 %. Fluka; Analytical; Chemical Abstracts Service (CAS)-No: 58969-01-0). Cell wall polysaccharides were stained with 0.1 % calcofluor white M2R (Sigma; CAS-No: 4404-43-7). To visualize lignin using brightfield microscopy, phloroglucinol staining was performed as described (9). Samples were incubated in 2 % phloroglucinol w/v in 95 % ethanol for 10 min, then staining solution was removed and samples were washed with 10 N HCl for 1 min. Samples were mounted and imaged in 1 N HCl solution.

**4-hydroxyindole histochemistry**

For *in tissue* laccase activity assays, transverse sections of mature fruit were incubated for 1 hour in 100 µM 4-hydroxyindole (Tokyo Chemical Industry; CAS-No: 2380-94-1) in the dark with shaking. Sections were washed in water two times before imaging. Laccase activity was detected as a dark blue precipitate on cell walls. To inhibit protein activity, sections were heated to 100°C in a heat block for 12 min, then incubated in 4-hydroxyindole. Sections were treated with 5 mM $CuSO_4$ with shaking for 30 min, washed three times with water and then incubated in 4-hydroxyindole.

**Quantification of Cu concentration in fruit**

Mature fruit were harvested and dried overnight in a vacuum desiccator Alpha 1-4 LSCplus (Martin Christ$^{TM}$). 5 to 7 fruit were pooled for each biological replicate. Dry weight was assessed and samples were digested with 1 ml of 67 % nitric acid in a 98 °C water bath. Digested samples were diluted by adding 9 ml of milliQ water to reach a final acid concentration of < 5 %. Samples were filtered using CellTrics disposable filters of 20 µm diameter (SYSMEX). Copper content was quantified by Inductively Coupled Plasma Mass Spectrometry using an Agilent 7700 ICP-MS (Agilent Technologies, Santa Clara, CA, United States) at the Biocenter Mass Spectrometry Platform, University of Cologne. Measurements of Cu ion content were normalized by the dilution factor of the sample and expressed as mg/kg dry biomass of mature fruit. One outlier data point (30.84 mg/kg) was removed in Fig. S2F as the difference of this value to the mean (18.71) was more than 10 times greater than the standard deviation.

**Lignin quantification and monomer analysis**

Lignin analyses were carried out as described (16). In brief, each biological replicate consists of 16 valves from 8 fruit at stage 17b pooled from 2 different plants. Samples were freeze-dried overnight. Alcohol insoluble residue (AIR) was prepared from the samples and enzymatically destarched. The lignin content was determined by quantifying the acetyl-bromide soluble lignin using spectrophotometry. The lignin composition was determined by thioacidolysis and TMS derivatization of the samples. The derivatives were then subjected to gas-chromatographic separation equipped with a SLB-5m capillary column (Supelco) and quantified using a quadrupole mass spectrometer (5977A Series, Agilent) compared to known standards.

**Laccase enrichment**

Laccase purification was performed as in (17) with several modifications. 10.5 g of mature wild-type *C. hirsuta* fruits were collected and immediately frozen in liquid nitrogen, stored at -80 °C and homogenized using a mortar and pestle in liquid nitrogen. 45 ml of extraction buffer (50 mM Tris-HCl, pH 8, 1.5 mM $CaCl_2$, 1M NaCl, 0.1 % w/v PVP90, 1 cOmplete ULTRA tablet [Roche 06538282001], 1 mM AEBSF, 2 mM DTT) were added to the homogenized tissue and mixed for 1 hour in a programmable rotator-mixer RM Multi-1 (Starlab) in a cold room. The homogenate was centrifuged 4 times, transferring the supernatant to a new tube after each centrifugation step (5 min, 3,000 g, 8 °C; 5 min, 3,000 g, 8 °C; 5 min, 13,000 g, 4 °C; 5 min, 15,000 g, 4 °C). The extraction was paper-filtered and then dialyzed using a SnakeSkin$^{TM}$ Dialysis Tubing (Thermo Scientific) three times in freshly prepared equilibration buffer (50 mM Tris, 0.5 mM NaCl pH 7.4), once in 2 l overnight and two times in 1 l buffer for one hour each. The dialysate was centrifuged at 15,000 g for 20 min at 4°C and the supernatant was subjected to affinity chromatography using Concanavalin A (Con-A) Agarose beads (C7555-5ML, Sigma) in an Econo-Column (7371512, Bio-Rad) with Econo-Column Flow Adaptor (7380016, Bio-Rad) and using a peristaltic pump (Rotarus Volume 50; Hirschmann). First, Con-A beads were pre-washed with 60 ml washing buffer (50 mM Tris, 0.5 M NaCl, pH 7.4) and equilibrated with equilibration buffer (50 mM Tris, pH 7.5, 0.5 M NaCl, 1mM $MgCl_2$, 1 mM $CaCl_2$, 1 mM $MnCl_2 \cdot 4\ H_2O$). 17 ml supernatant were loaded on the column and then the column was washed with 50 ml of washing buffer. Column was eluted with Methyl α-D-mannopyranoside in three steps (3 ml each) with increasing concentration (100, 200 and 350 mM) and, afterwards, column was washed with 29 ml of washing buffer. The flow through was collected in 3 ml fractions. Five fractions with high protein concentration were pooled to use for *in gel* analysis of laccase activity.

***In gel* laccase activity assay**

25 µg of Con-A enriched laccases were run on a non-denaturing 10 % Mini-PROTEAN TGX gel (Bio-Rad). 25 ng of laccase from *Aspergillus sp*. (CAS-No: 804-15-3) were used as control. Gel was washed three times in water for a few seconds, then incubated in 5 mM 4-hydroxyindole (30 ml water + 300 µL of stock 500 mM 4-hydroxyindole dissolved in ethanol) with gentle shaking for 30 min before imaging.

**RNA sequencing**

Fruit valves of wild type and *spl7-1* mutant were dissected from fruit at stage 17 (at the onset of SCW lignification in end*b* cells). Three biological replicates per genotype were analyzed. RNA was extracted from approximately 15 pooled valves for each biological replicate using the Spectrum Total RNA kit (Sigma-Aldrich). cDNA synthesis was performed using SuperScript III Reverse Transcriptase (Invitrogen). TruSeq libraries were prepared and sequenced using the HiSeq2500 Illumina platform at the MPIPZ Genome Centre. 20,000,000 paired-end reads of 100 bp were requested for each library. Paired-end reads were aligned to the *C. hirsuta* reference genome (18) using TopHat2 (19) and raw read counts were quantified with HTSeq v0.5.4 (20). Principal component analysis and hierarchical clustering were used to visualize the overall similarity between samples and assess RNA-seq data quality. The software package DESeq from Bioconductor (21) was used to test for differential gene expression between genotypes

**Quantitative RT-PCR**

Gene expression was quantified in stage mature fruits of *C. hirsuta* genotypes that varied for *SPL7* copy number. Tissue was collected and flash-frozen in liquid nitrogen. Three biological samples, consisting of two pooled fruits each, were analysed per genotype. Total RNA was extracted using the Spectrum Total RNA kit (Sigma-Aldrich). 550 ng of RNA was used for cDNA synthesis using the SuperScriptIV kit (Invitrogen). *SPL7* was amplified using a QuantStudio5 thermocycler (Applied Biosystems) from the cDNA in a qPCR reaction using Power SYBR Green Supermix (Applied Biosystems). Expression levels were normalized against the housekeeping gene *TIP41* (CARHR242510). Primers used are listed in Table S2.

**Statistical Analyses**

Statistical analyses were done with R Statistical Software (22). For parametric statistical analysis, one-way ANOVA and Tukey's test were used. For nonparametric statistical analysis, Kruskal-Wallis and Fisher's least significant difference test or Dunn's test as post hoc analysis were used. Binary comparisons were performed using Student's t test.

**Table S1. Oligonucleotides used in this study.**

| Primer name | Sequence (5´ - 3´) | Experiment |
|---|---|---|
| pSPL7fullF | GGTCTCAACCTTAACCACCACCAAAGGTAACAA | Cloning of pSPL7 |
| pSPL7fullR | GGTCTCATGTTCTGAGTCTTCTTCAATTCATAAATTTC | Cloning of pSPL7 |
| cdsSPL7_F | GGTCTCAGGCTCCATGTCTTCTCTGTCGCAATCC | Cloning of SPL7 CDS |
| cdsSPL7stopR | GGTCTCACTGATTAAATTCTGCGTATCAATCTCATC | Cloning of SPL7 CDS |
| SBPcdsSPL7_R | GGTCTCACTGACTGGTCAACAGAGCATGTATTATCTT | Cloning of SBP domain |
| pSPL7_attB4_F | GGGGACAACTTTGTATAGAAAAGTTGCTTAACCACCAC CAAAGGTA | Cloning of pSPL7 |
| pSPL7_attB1r_R | GGGGACTGCTTTTTTGTACAAACTTGCCTGAGTCTTCTT CAATTCA | Cloning of pSPL7 |
| cdsSPL7_attB1_F | GGGGACAAGTTTGTACAAAAAAGCAGGCTTAATGTCTT CTCTGTCGCAA | Cloning of SPL7 CDS |
| SPL7NoStopB2_R | GGGGACCACTTTGTACAAGAAAGCTGGGTAAATTCTGC GTATCAATCTC | Cloning of SPL7 CDS |
| proLAC4_F | GGTCTCAACCTCAACATTGTAATGGACTTGAATCT | Cloning pLAC4 |
| proLAC4_R | GGTCTCATGTTCCCTCTAGCTCTCTCTATTCTCTCT | Cloning pLAC4 |
| proLAC11_F1 | GGTCTCAACCTATAACTTTTCAAAGTCTCGACCTTTTT | Cloning pLAC11 |
| proLAC11_R1 | GGTCTCAGTCCACAAGATTTGAAAATGAAGCAT | Cloning pLAC11 |
| proLAC11_F2 | GGTCTCTGGACTCAATTATCGATAAATCTACACAT | Cloning pLAC11 |
| proLAC11_R2 | GGTCTCATGTTTTCCGGTTCAATCTTCCGGTT | Cloning pLAC11 |
| proLAC17_F | GGTCTCAACCTGAAACAGATTTTGATTCTCCTCAA | Cloning pLAC17 |
| proLAC17_R | GGTCTCATGTTTTTAAGTGAGCTTGGAACCCG | Cloning pLAC17 |
| cdsLAC4_F | GGTCTCAGGCTCCATGGGATCTCATATGGTTTGG | Cloning LAC4 CDS |
| cdsLAC4NoStopR | GGTCTCACTGAACATTTGGGAAGATCCTTAGGC | Cloning LAC4 CDS |
| cdsLAC11_F | GGTCTCAGGCTCCATGAAGATCATCCGAGTCCCG | Cloning LAC11 CDS |
| cdsLAC11NoStop | GGTCTCACTGAGCACGACGGATAGTCTTTAGG | Cloning LAC11 CDS |

| | | |
|---|---|---|
| cdsLAC17_F | GGTCTCAGGCTCCATGGCGTTTCAGCTTCTCCT | Cloning LAC17 CDS |
| cdsLAC17NoStop | GGTCTCACTGAGCATTTGGGCAAGTCTGC | Cloning LAC17 CDS |
| proPER49_F | AACAGGTCTCAACCTAGACTTGGACTTTGTCTACGC | Cloning of pPER49 |
| proPER49_R | AACAGGTCTCATGTTTACTTTCAACAAGAAGATGAGGAA | Cloning of pPER49 |
| proPER64_F | AACAGGTCTCAACCTTGCCATTAAGAGGAGGTTACAA | Cloning of pPER64 |
| proPER64_R | AACAGGTCTCATGTTTTCAGCAAATGTTTCGAAATC | Cloning of pPER64 |
| proPER66_F | AACAGGTCTCAACCTTATTTAACGTGTAATGGGGTTTTT | Cloning of pPER66 |
| proPER66_R | AACAGGTCTCATGTTTTTGATGATGCAGAAGAAGAAGC | Cloning of pPER66 |
| cdsPER49_F | AACAGGTCTCAGGCTCCATGGCAAGACTCACTAGCTTTC | Cloning of PER49 CDS |
| cdsPER49NoStop | AACAGGTCTCACTGAAGAGTTTATCTTCCTGCAATTCTTC | Cloning of PER49 CDS |
| cdsPER64_F | AACAGGTCTCAGGCTCCATGAATGCAAATATACTGATCAATCTC | Cloning of PER64 CDS |
| cdsPER64NoStop | AACAGGTCTCACTGAGCGAACCCTCCTGCAATTA | Cloning of PER64 CDS |
| cdsPER66_F | AACAGGTCTCAGGCTCCATGTCATTCTCGAAAGGACTCA | Cloning of PER66 CDS |
| cdsPER66NoStop | AACAGGTCTCACTGAGTTGATGAAGCGAGTTTTAA | Cloning of PER66 CDS |
| qPCR_TIP41_F | GATGGTGTGCTTATGAGATTGAGAG | qRT_PCR TIP41 (CARHR242510) |
| qPCR_TIP41_R | TCAACTGGATACCCTTTCGCA | qRT_PCR TIP41 (CARHR242510) |
| qPCR_SPL7_F2 | TGAAGCTCAGCCAGATGAAGG | qRT_PCR SPL7 |
| qPCR_SPL7_R2 | CGTGGGAACTCTGCTGGATT | qRT_PCR SPL7 |
| INDEL_16810437_F | CAATTCATAGTTGTGGATGTCTTCTC | Mapping *lig1* INDEL16810437 (Ox + 53 bp) |
| INDEL_16810437_R | CGGGCGGACACAATAGTTTA | |
| SNP_16824303_F | CACATCTTTAAACAAATGCCGATCTA | Mapping *lig1* SNP16824303 (dCAPS/*Xba*I cuts Nz) |
| SNP_16824303_R | ACAACCGCGCTTTTATTCAT | |
| INDEL_17317339_F | TTGTTGTGAAACTTATTTTGTGGTC | |

| | | |
|---|---|---|
| INDEL_17317339_R | AATAACCGCCATTTTAACAATTC | Mapping *lig1* INDEL17317339 (Ox + 21 bp) |
| SNP_17676369_F | TGTAATGGAAGTGGGAACAGAAGTA | Mapping *lig1* SNP17676369 (dCAPS/*Sca*I cuts Ox) |
| SNP_17676369_R | CCATAATCTTTATTAGCACATCGAG | |
| INDEL_17882576_F | TCCACGTGGGGTAAAATAATG | Mapping *lig1* INDEL17882576 (Ox + 65 bp) |
| INDEL_17882576_R | CCTGAATCTGGTGTCGTAGC | |
| SNP_18274972_F | TCTTCTTCCTGATTTCTACATTCTA | Mapping *lig1* SNP18274972 (dCAPS/*Xba*I cuts Ox) |
| SNP_18274972_R | TCCGTATTAATCAAAATCGAAGC | |
| INDEL_18607525_F | CACAAGTCGTCGTCTCTTGC | Mapping *lig1* INDEL18607525 (Ox + 31 bp) |
| INDEL_18607525_R | CCCGTTTCCATGAATCTGTT | |

**Table S2. sgRNA protospacers for CRISPR/Cas9 mutagenesis used in this study.**

| sgRNA name | sgRNA sequence (5´ - 3´) |
|---|---|
| *SPL7* sgRNA-1 | ATCCCCATCGGCGCCTGAGA***TGG*** |
| *SPL7* sgRNA-2 | GTTCGTCAGCGGCGAAGTCA***AGG*** |
| *LAC4* sgRNA-1 | GCCAACGTGGAACGCTCTGG***TGG*** |
| *LAC4* sgRNA-2 | CACATGATTAACGGACACCC***TGG*** |
| *LAC11* sgRNA-1 | CCGGACAACGCGGGACCCTC***TGG*** |
| *LAC11* sgRNA-2 | TTTCTACTACGGTCATGTTG***TGG*** |
| *LAC17* sgRNA-1 | CCGGCAATTACGGAGTGGTT***GGG*** |
| *LAC17* sgRNA-2 | GAGCAGTTGTATAACGGACC***AGG*** |
| *PER66* sgRNA-1 | AGGATACGAGCAGGCACTTT***GGG*** |
| *PER66* sgRNA-2 | TCAGCACAAGAAACGGTACG***AGG*** |

### Supplementary figures

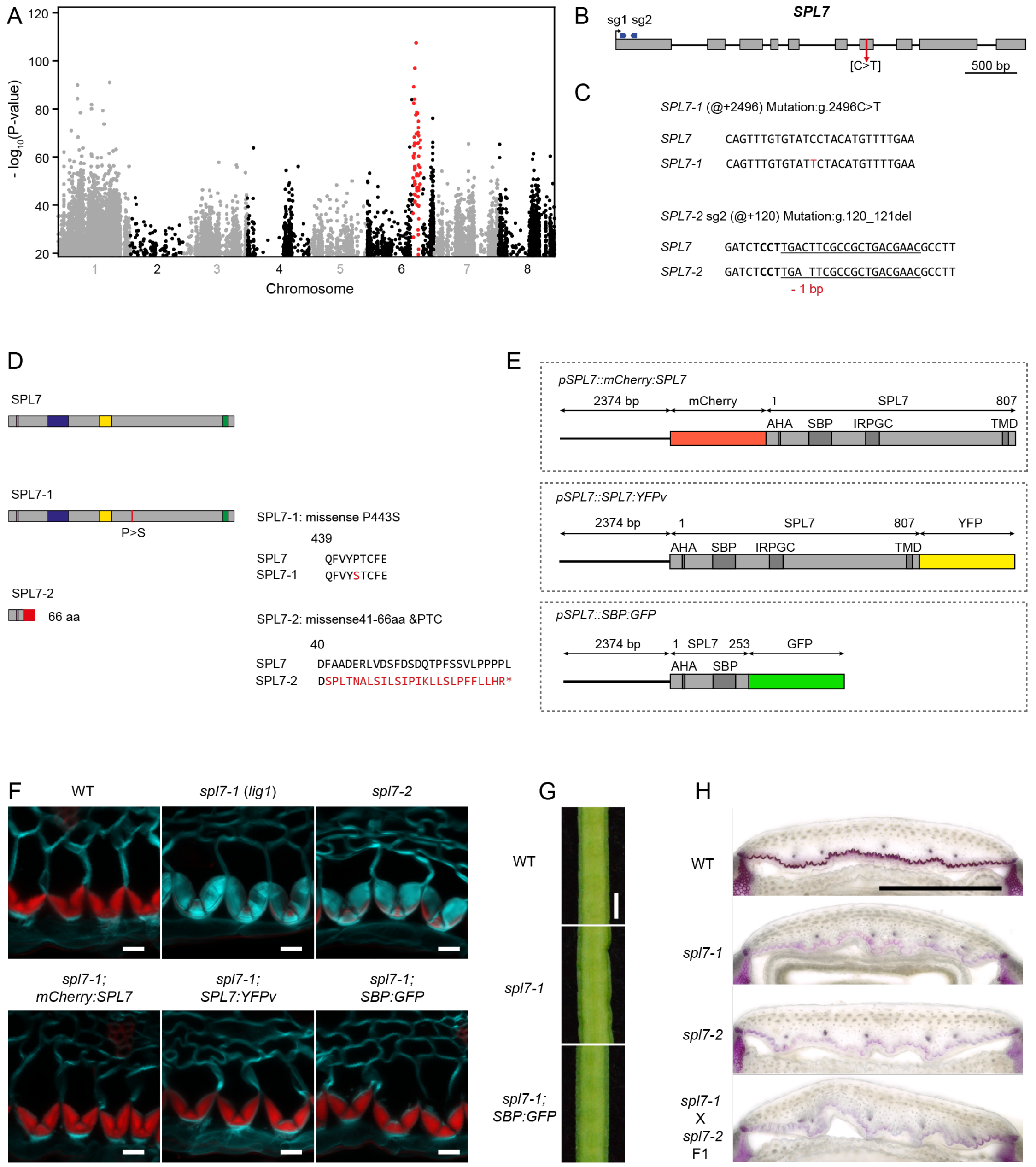


**Fig. S1. Characterization of *C. hirsuta spl7* mutant alleles.** (*A*) Manhattan plot of -log10(*P*-value) (Fisher's exact test) for SNPs identified genome-wide in *C. hirsuta lig1* (*spl7-1*) mapping-by-sequencing data. SNPs in the region with overall low *P*-values in chromosome 6 are shown in red. Only SNPs with -log10(P-value) > 18.9 were included in this analysis. (*B*) Schematic representation of *SPL7* gene showing positions of *spl7-1* (*lig1*) mutation and CRISPR sgRNAs used to generate *spl7-2* allele. Gray boxes: coding DNA sequences, black lines: introns, red arrow: *spl7-1* mutation (C to T substitution), blue arrows: sequences targeted by CRISPR sgRNAs. (*C*) Description of *spl7* mutant alleles. @+ symbol on wild-type sequence indicates position of mutation counting from the start codon. Nomenclature as in (23). (*Top*) EMS-derived *spl7-1* (*lig1*) allele. Nucleotide substitution indicated in red.

(*Bottom*) *spl7-2* allele. Underlined sequence: CRISPR sgRNA, bold text: protospacer adjacent motif (PAM). Mutation indicated in red. (*D*) Putative translational products of *spl7* mutant alleles. (*Left*) Schematic representation of the protein encoded by the wild-type and mutant alleles of *SPL7*. Pink box: AHA-like transcriptional activator, blue box: SQUAMOSA-PROMOTER BINDING DOMAIN (SBP), yellow box: IRPGC domain, green box: transmembrane domain (TMD). Red box indicates putative peptide from missense translation in *spl7-2* allele. Mutant *spl7-2* allele is represented up to the premature termination codon (PTC). (*Right*) Predicted peptide sequence encoded by the wild-type and *spl7* mutant alleles. Red characters indicate amino acid residues that result from missense translation in the mutants. Above, a description of the mutant protein is provided indicating the number of aa residues resulting from missense translation and the PTC. For instance, "missense41-66aa &PTC" indicates that aa residues 41 to 66 result from missense translation, followed by a PTC. (*E*) Schematic representation of SPL7 translational fusions used for complementation of *spl7-1* mutant. Black line: promoter sequence, light grey box: coding DNA sequence, dark grey boxes: conserved domains of SPL7 protein. Numbers above grey boxes indicate length of the SPL7 peptide encoded by the transgene. Number above black lines indicates the length, in bp, of the *SPL7* promoter amplified to generate the constructs. (*F*) Lignin, stained red with Basic Fuchsin, and cell walls stained cyan with Calcofluor White, in end*b* cells of wild type, *spl7-1*, *spl7-2* and the following complementation lines: *spl7-1; mCherry:SPL7*, *spl7-1; SPL7:YFPv* and *spl7-1; SBP:GFP*. Confocal micrographs show z-axis sum projections. (G) Close-up views of the fruit margin in wild type, *spl7-1* and *spl7-1; SBP:GFP*. (*H*) Lignin stained with phloroglucinol in cross sections of wild-type, *spl7-1*, *spl7-2* and *spl7-1*/*spl7-2* trans-heterozygote fruit. Plants were supplemented with 0.5 mM $CuSO_4$ to ensure plant growth and fruit development in *spl7* mutants (*F*, *G*, *H*). Scale bars: 10 µm (*F*), 1 mm (*G*), 500 µm (*H*).

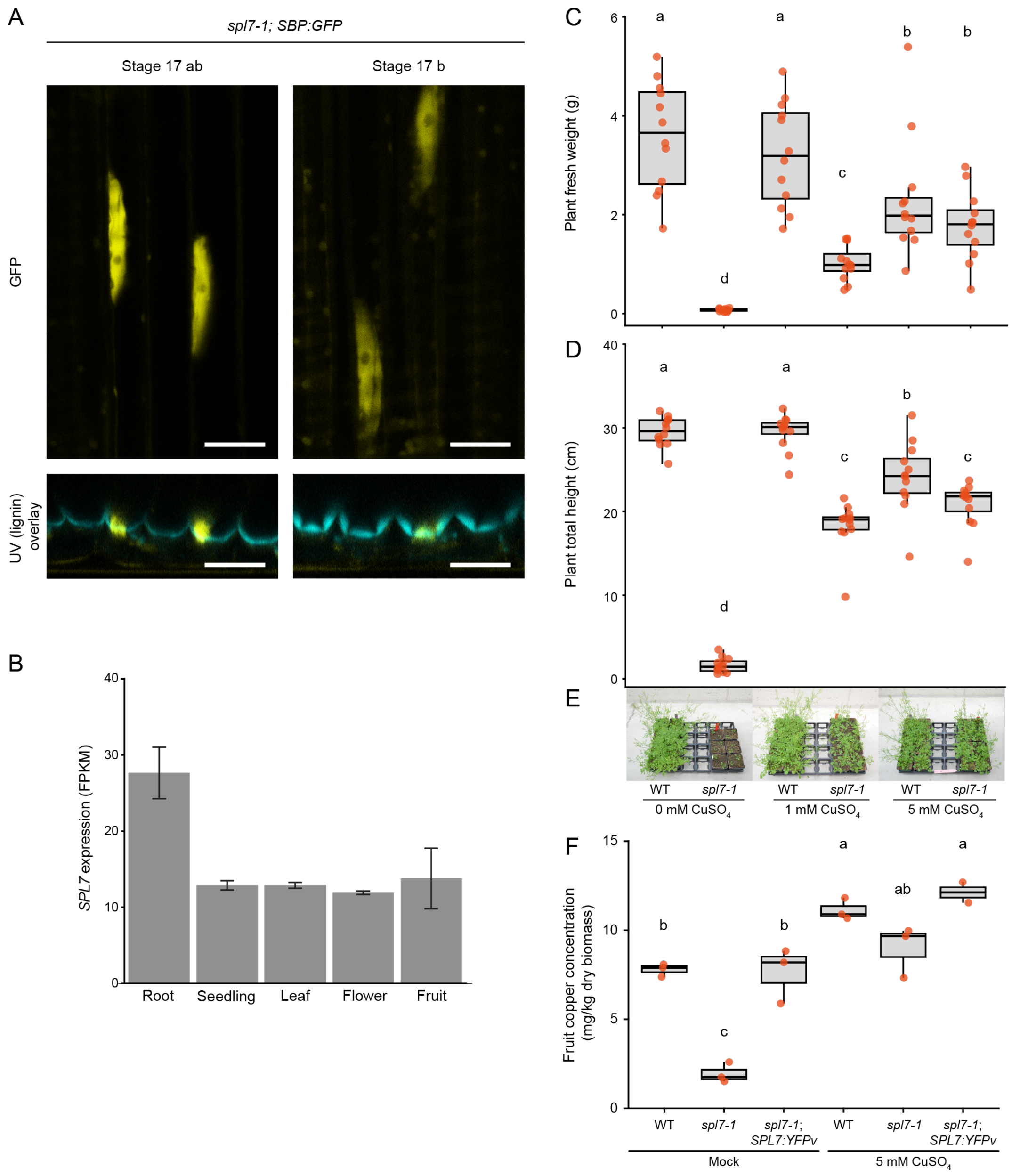


**Fig. S2. Copper supplementation rescues *C. hirsuta spl7* mutant phenotypes.** (*A*) SBP:GFP localization (yellow) in end*b* cell nuclei of stage 17ab and 17b fruit valves from *spl7-1; pSPL7::SBP:GFP* plants, imaged *en face* (top, z-axis sum projections) and shown together with lignin autofluorescence (cyan) in transverse optical sections (bottom, sum projections of several optical transverse sections). (*B*) *SPL7* expression in *C. hirsuta* tissues, taken from several RNAseq experiments and indicated as Fragments Per Kilobase Million (FPKM). Root: roots from 14 days old seedlings grown in ½ MS media under SD conditions (n = 3 biological replicates), seedling: 14-days-old seedlings grown in ½ MS media under SD conditions (n = 3 biological replicates), leaf: developing leaves of plants grown in LD conditions (n = 3 biological replicates). Flower: stage 9 flowers from plants in LD conditions (n = 2 biological replicates), fruit: stage 16 fruit from plants in LD conditions (n = 2 biological replicates) (18).

(*C-E*) Boxplots of fresh weight (*C)* and height (*D*) of aerial parts from 7-week-old wild-type and *spl7-1* plants in response to mock (water), 1 mM or 5 mM $CuSO_4$ treatments (regular 20 ml applications). Plots show median (thick black line), n = 12 plants per genotype per treatment (red dots), different letters denote statistical significance at $P < 0.05$ using Kruskal-Wallis test and Fisher's least significant difference as post hoc analysis; 7-week-old plants shown in (*E*). (*F*) Boxplot of Cu concentration (mg/kg dry biomass) in mature fruits of wild type, *spl7-1* and *spl7-1; SPL7:YFPv* plants grown on soil in response to mock (water) or 5 mM $CuSO_4$ treatments (20 ml applications); n = 3 biological replicates per condition (red dots) where each replicate contains 4 fruit from one plant. Scale bars: 20 µm (*A*).

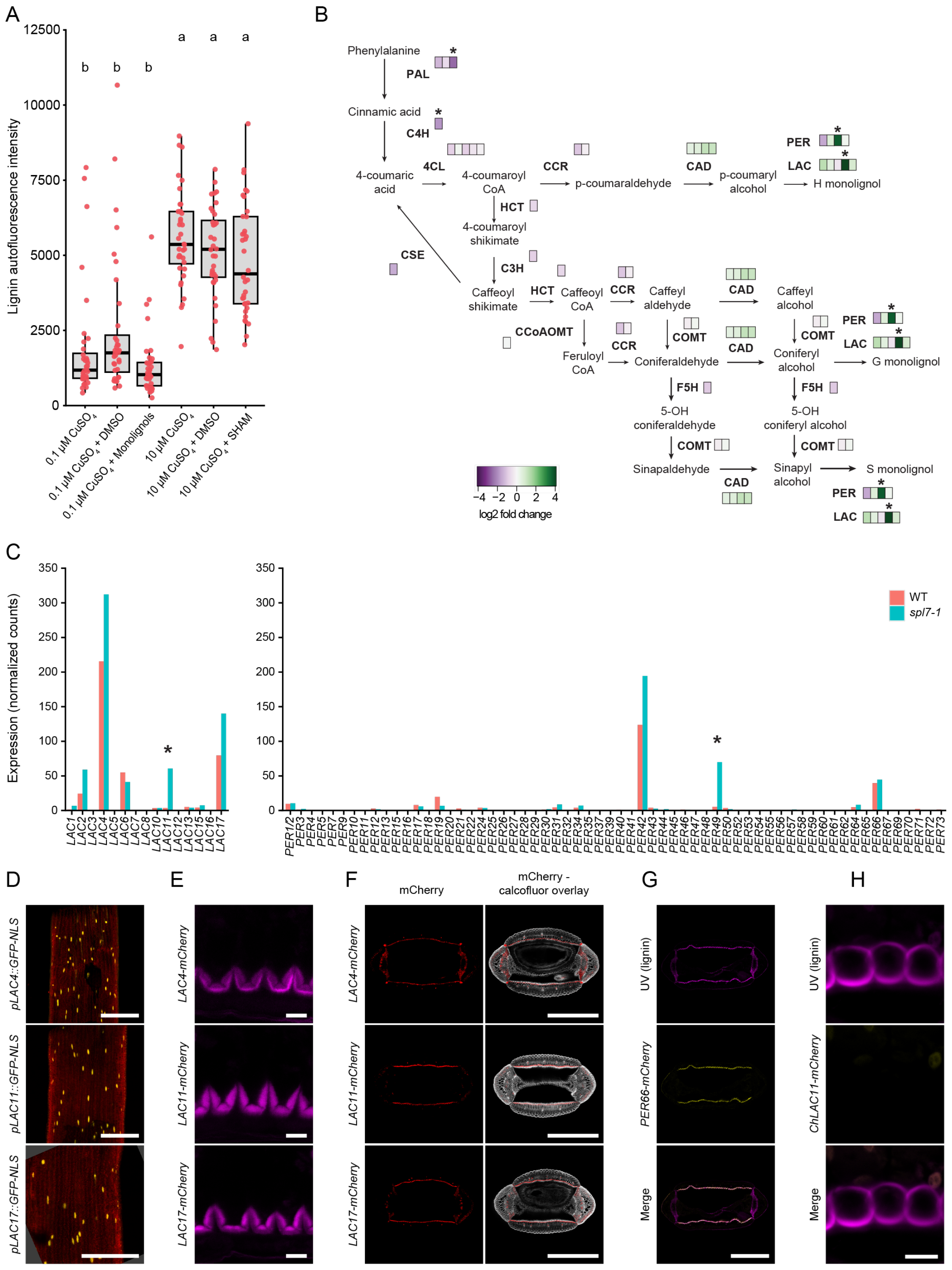


**Fig. S3. Laccase and peroxidase expression in *C. hirsuta* fruit.** (*A*) Boxplot of lignin autofluorescence in endb cells of *spl7-1* fruit grown in media containing either 0.1 μM $CuSO_4$ and treated with monolignols (100 μM coniferyl and sinapyl alcohols) vs mock (DMSO), or 10 μM $CuSO_4$ and treated with peroxidase inhibitor (150 μM SHAM) vs mock (DMSO); n = 36 endb cell regions per treatment (red dots) from 9 similar-stage fruits from multiple plants. Plants were grown on soil and supplemented with 0.5 mM $CuSO_4$ to ensure growth and development before fruit were excised.

Different letters denote statistical significance at $P < 0.05$ using Kruskal-Wallis and Dunn's test as post hoc analysis. (*B*) Differential expression of lignin biosynthesis genes between fruit valves of wild type and *spl7-1* (also see Fig. 3A main text and Dataset S1). Values of log2-fold change are represented with a heatmap color scale and mapped to a diagram of the lignin biosynthesis pathway. On each biosynthetic reaction, each rectangle represents a gene encoding for a protein potentially involved in lignin biosynthesis. Only genes with a mean value of wild type and *spl7-1* normalized counts >10 are presented. Asterisks denote statistically significant differences in gene expression between wild type and *spl7-1* mutant at adjusted *P*-value < 0.05. Genes can encode proteins involved in several different steps of the pathways leading to the synthesis of the different monolignols, therefore, genes and log2-fold change values are indicated in all steps where they are potentially involved. (*C*) Barplots of gene expression of laccases (*Left*) and peroxidases (*Right*) in *C. hirsuta* wild-type and *spl7-1* fruit valves shown as normalized counts (also see Fig. 3A main text and Dataset S1). Asterisks denote statistical significance in gene expression between genotypes at adjusted *P*-value < 0.05. (*D*) Expression of *pLAC4::GFPnls*, *pLAC11::GFPnls* and *pLAC17::GFPnls* (yellow) promoter reporters in end*b* cells of *C. hirsuta* mature fruit valves, imaged *en face* and shown together with chlorophyll autofluorescence (red). (*E*) *C. hirsuta* LAC4, LAC11 and LAC17 protein fusions (magenta, *pLAC::LAC:mCherry*) localize to *C. hirsuta* end*b* SCWs. These same images are also shown in overlay with calcofluor in Fig. 3B main text. (*F*) Localization of *C. hirsuta* LAC4, LAC11 and LAC17 protein fusions (red, *pLAC::LAC:mCherry*) in cross sections of *C. hirsuta* mature fruit. (*Left*) mCherry, red. (*Right*) Overlay of mCherry and calcofluor white (gray). (*G*) Localization of *C. hirsuta* PER66:mCherry protein fusion (yellow) in cross section of *C. hirsuta* mature fruit, lignin shown as UV autofluorescence (magenta). (*H*) *C. hirsuta* LAC11:mCherry protein fusion (yellow) does not accumulate in SCWs of end*b* cells of *A. thaliana* fruit, lignin shown as UV autofluorescence (magenta). Confocal micrographs show z-axis sum projections (*D-H*). Abbreviations: PAL: phenylalanine ammonialyase, C4H: cinnamate 4-hydroxylase, 4CL: 4-hydroxy cinnamoyl CoA ligase, CSE: Caffeoyl shikimate esterase, HCT: hydroxycinnamoyl-CoA shikimate/quinate hydroxycinnamoyl transferase, C3′H: p-coumaroyl shikimate/quinate 3′-hydroxylase, CCR: (hydroxy) cinnamoyl CoA reductase, CAD: (hydroxy) cinnamyl alcohol dehydrogenase, CCoAOMT: caffeoyl CoA O-methyl transferase, F5H: ferulic acid/coniferaldehyde/coniferyl alcohol 5-hydroxylase, COMT: caffeic acid/5-hydroxyferulic acid O-methyltransferase, PER: peroxidase, LAC: laccase. Scale bars: 500 µm (*D*, *F*, *G*), 10 µm (*E*, *H*).

A

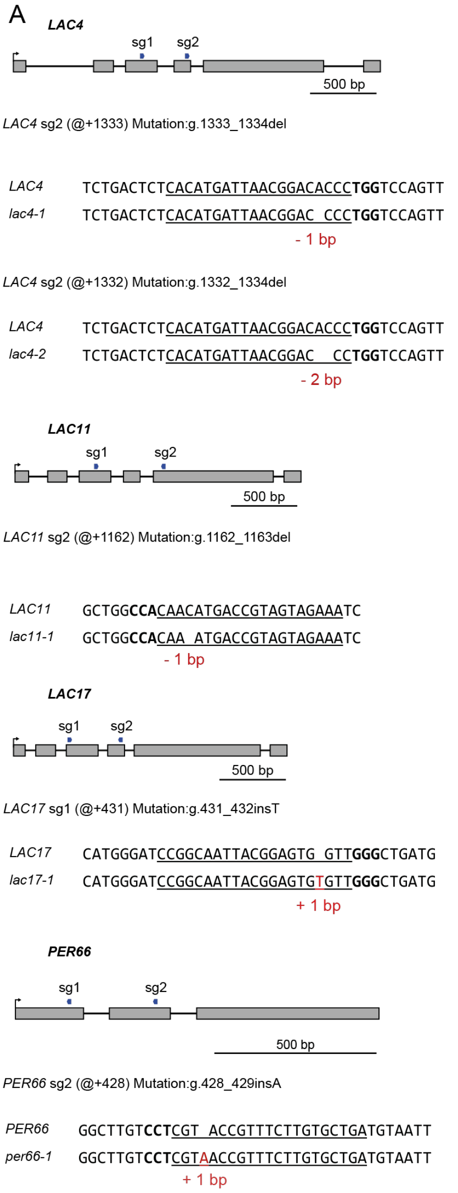


B

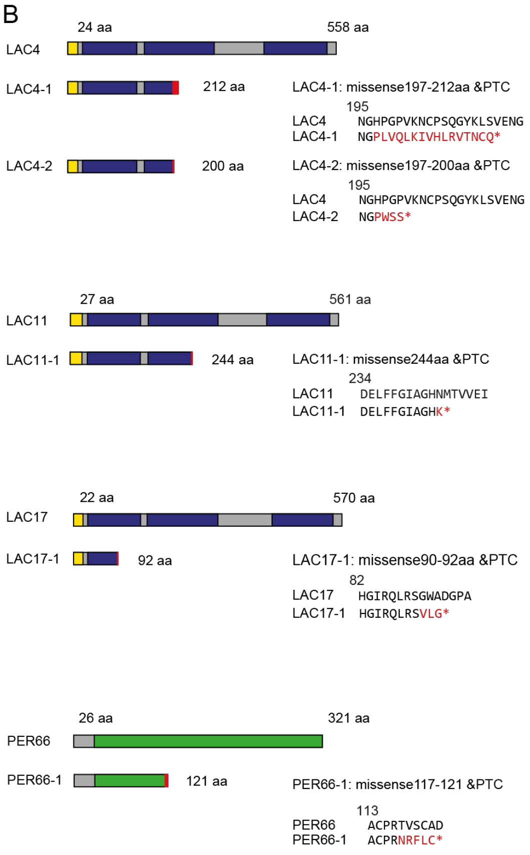


C

| | Allele | | | |
|---|---|---|---|---|
| line | *lac4* | *lac11* | *lac17* | *per66* |
| *lac4* | 1 | | | |
| *lac11* | | 1 | | |
| *lac17* | | | 1 | |
| *per66* | | | | 1 |
| *lac4 lac11* | 2 | 1 | | |
| *lac4 lac17* | 1 | | 1 | |
| *lac11 lac17* | | 1 | 1 | |
| *lac4/+ lac11 lac17* | 1-/+ | 1 | 1 | |
| *lac4 lac11/- lac17* | 1 | 1-/+ | 1 | |
| *lac4 lac11/- lac17 per66* | 1 | 1-/+ | 1 | 1 |

**Fig. S4. Characterization of CRISPR/Cas9 alleles of *LAC4*, *LAC11*, *LAC17* and *PER66*.** (*A*) Schematic representation of *LAC4*, *LAC11*, *LAC17* and *PER66* genes and CRISPR sgRNAs used in this study. Gray boxes: coding DNA sequences, black lines: introns, blue arrows: sequences targeted by CRISPR sgRNAs. Description of *lac4*, *lac11, lac17* and *per66* mutant alleles. @+ symbol on wild-type sequence indicates position of mutation counting from the start codon. Nomenclature as in (23). For example, g1333_1334del indicates a deletion of one nucleotide. Mutations are indicated in red. Underlined sequence: CRISPR sgRNA, bold text: PAM sequence. (*B*) Putative translational products of the *lac4*, *lac11, lac17* and *per66* mutant alleles generated by CRISPR-Cas9. (*Left*) Schematic representation of the proteins encoded by wild-type and mutant alleles. Blue box: multicopper oxidase domain, yellow box: signal peptide, green box: peroxidase domain. Red rectangle indicates putative peptide from missense translation in mutant alleles. Mutant alleles are represented up to the premature termination codon (PTC). (*Right*) Predicted peptide sequence encoded by wild-type and mutant alleles. Red characters indicate amino acid residues that result from missense translation. Description of each mutant protein, indicating the number of aa residues resulting from missense translation and the PTC. For example, "missense197-121aa &PTC" indicates that aa residues 197 to 121 result from missense translation, followed by a PTC. (*C*) Summary of *lac4*, *lac11, lac17* and *per66* alleles in the genotypes analyzed in this study. Alleles are indicated by number, note that a single allele is used in all cases except two *lac4* alleles; +/- indicates heterozygous genotype.

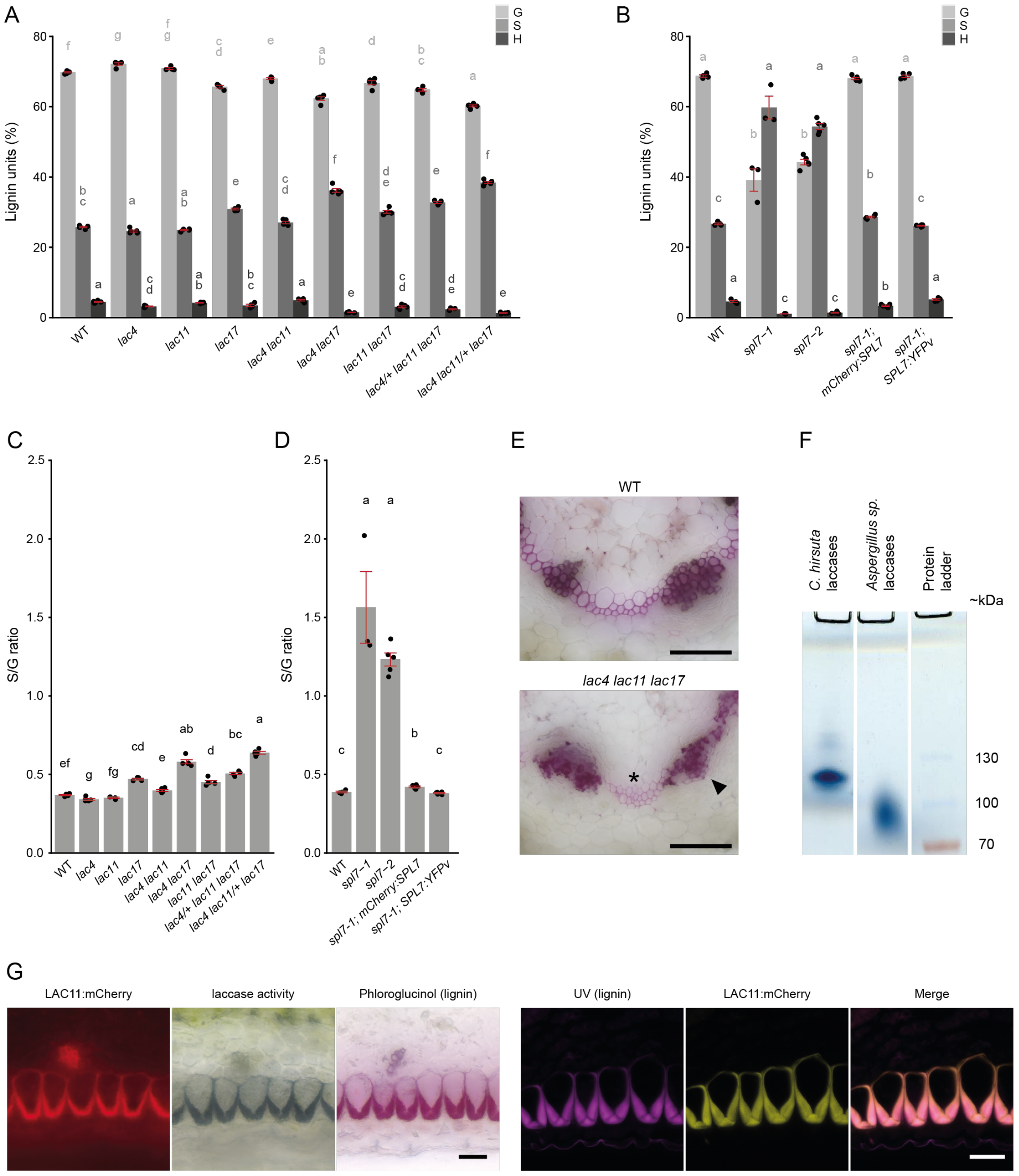


**Fig. S5. Comparison of lignin composition between *spl7* and *lac4, lac11* and *lac17* alleles.** (*A-D*) Barplots of lignin monomer composition obtained by thioacidolysis (*A-B*), and S/G monomer ratio (*C-D*), in fruit valves of wild type, *lac4, lac11* and *lac17* allelic combinations (*A, C*) and wild type, *spl7-1, spl7-2, spl7-1 mCherry:SPL7* and *spl7-1 SPL7:YFPv* complementation lines (*B, D*). Y-axes indicate the contribution of each monolignol to total lignin as % (*A-B*), and S/G monomer ratio (*C-D*). G: coniferyl alcohol, S: sinapyl alcohol, H: *p*-coumaryl alcohol. Plots show mean, error bars indicate S.E.M., n = 3-5 replicates per sample (black dots), different letters denote statistical significance at $P < 0.05$ using Kruskal-Wallis and Fisher's least significant difference as post hoc analysis. (*C-D*) Barplots of S/G monomer ratio in the same samples as in (*A-B*). (*E*) Lignin, stained purple with phloroglucinol, in cross sections of wild-type and *lac4 11 17* triple mutant stems. Arrow head indicates collapsed xylem vessels. Asterisk indicates reduced lignin deposition in fibers. (*F*) *In gel* laccase activity assay of Con-A enriched proteins from wild-type mature fruit, visualized as precipitation of oxidized 4-hydroxyindole products. Laccases from *Aspergillus sp*. are used as positive control. (*G*) Extended dexamethasone induction (see methods) of *lac4 11 17 pLAC11::LhGR>>LAC11:mCherry* plants grown in LD conditions. End*b* cells showing, from left to right, ectopic LAC11:mCherry localization (red) and laccase activity (black oxidized 4-hydroxyindole precipitate) in the same cells, ectopic lignin (stained with phloroglucinol), ectopic LAC11:mCherry localization (yellow) and lignin autofluorescence (magenta) imaged by CLSM. Images show z-axis extended depth of field projections and CLSMs are sum projection images. Scale bars: 100 µm (*E*), 20 µm (G).
